# Hepatocyte nuclear factor 4a and glucocorticoid receptor coordinately regulate lipid metabolism in mice fed a high-fat-high-sugar diet

**DOI:** 10.1101/2021.02.06.427306

**Authors:** Hong Lu, Xiaohong Lei, Shangdong Guo, Rebecca Winkler, Savio John, Devendra Kumar, Wenkuan Li, Yazen Alnouti

**Author notes:** To whom correspondence should be addressed: Hong Lu: Department of Pharmacology, SUNY Upstate Medical University, Syracuse NY 13210.

## Abstract

Hepatocyte nuclear factor 4α (HNF4α) and glucocorticoid receptor (GR), master regulators of liver metabolism, are down-regulated in fatty liver diseases. The present study was aimed to elucidate the role of down-regulation of HNF4α and GR in fatty liver and hyperlipidemia. Adult mice with liver-specific heterozygote and knockout (knockout) of HNF4α were fed a low-fat diet (LFD) or a high-fat-high-sugar diet (HFHS) for 15 days. Compared to LFD-fed mice, HFHS-fed wildtype mice had hepatic induction of lipid catabolic genes and down-regulation of lipogenic genes. Compared to HFHS-fed wildtype mice, HNF4α heterozygote mice had down-regulation of lipid catabolic genes, induction of lipogenic genes, and increased hepatic and blood levels of lipids, whereas HNF4α knockout mice had mild hypolipidemia, down-regulation of lipid-efflux genes, but induction of genes for uptake/storage of lipids. Sterol-regulatory-element-binding protein-1c (SREBP-1C), a master lipogenic regulator, was induced in HFHS-fed HNF4α heterozygote mice. In reporter assays, HNF4α potently inhibited the transactivation of mouse and human SREBP-1C promoter by liver X receptor. Surprisingly, nuclear GR proteins were gene-dosage-dependently decreased in HNF4α heterozygote and knockout mice. HFHS-fed mice with liver-specific knockout of GR had increased hepatic lipids and induction of SREBP-1C and PPARγ. In reporter assays, GR and HNF4α synergistically/additively induced lipid catabolic genes. Phosphorylation of AMP-activated protein kinase (AMPK), a key GR modulator, was dramatically decreased in HNF4α knockout mice. Thus, cooperative induction of lipid catabolic genes and suppression of lipogenic genes by HNF4α and GR, modulated by AMPK, may mediate the early resistance to HFHS-induced fatty liver and hyperlipidemia.

---

In modern society, the synergy between excessive psychological stress and overeating high-fat and high-sugar diets (HFHS) propels a pandemic of non-alcoholic fatty liver disease (NAFLD) which has now replaced viral hepatitis as the most common chronic liver disease (28). It is well known that high-fat-diet (HFD) does not cause fatty liver until after long-term feeding (65). A poorly understood knowledge gap is the molecular mechanism of early resistance of the liver to HFD/HFHS-induced steatosis and how the resistance is compromised over time; bridging this key knowledge gap will help discover novel preventive and therapeutic strategies for NAFLD.

Hepatocyte nuclear factor 4α (HNF4α) is a master regulator of liver metabolism and lipid homeostasis via crosstalk with diverse extracellular and intracellular signaling pathways to regulate hepatic nutrient metabolism (80). Hepatic HNF4α expression is markedly decreased in diabetes and NAFLD (73, 145, 146). Paradoxically, adult chow-fed mice with liver-specific knockout (knockout) of HNF4α have fatty liver but striking hypolipidemia and are protected from atherosclerosis (146). Thus, suppression of hepatic HNF4α has been proposed to prevent atherosclerosis (146). However, the marked hypolipidemia in chow-fed adult HNF4α knockout mice sharply contrasts with the hyperlipidemia in patients and animal models with NAFLD/non-alcoholic steatohepatitis (NASH) in which HNF4α is partially lost (7, 77), a condition that may be better mimicked in mice with hetero-deficiency of HNF4α. The most common cause of death among patients with NAFLD is cardiovascular disease (CAD) (75). In contrast to hyperlipidemia in NAFLD, hypolipidemia often occurs in patients with end-stage liver diseases, such as cirrhosis and liver cancer (105), in which the loss of functional HNF4α is more marked (34) and may be better mimicked by the use of HNF4α knockout mice. Thus, there is a key knowledge gap regarding the gene-dosage-dependent roles of partial and total loss of HNF4α in regulating hepatic lipid metabolism, circulating lipids, and CAD.

Because diet intake has a critical role in modulating hepatic lipid metabolism and circulating lipid profiles, the purpose of this study was to uncover how interactions between gene-dosage-dependent HNF4α deficiency and different diets may differentially alter hepatic gene expression and lipid metabolism. In previous studies (47, 82), the chow fed to adult HNF4α knockout mice was a low-fat diet (LFD). Similarly, we found that adult male HNF4α knockout mice fed a LFD had marked hypolipidemia which was attenuated by HFHS feeding. Interestingly, we found that although LFD-fed adult male mice with liver-specific heterozygote of HNF4α had normal blood lipid profiles, HFHS-fed HNF4α heterozygote mice had elevated hepatic and blood levels of cholesterol and triglycerides. Results from further mechanistic studies suggest that HNF4α has a key role in regulating hepatic lipid metabolism and circulating lipids by inducing lipid catabolic genes and suppressing lipogenic genes.

Consumption of HFHS “comfort foods” is associated with elevated circulating glucocorticoids (GCs) (24). Currently, the role of glucocorticoid receptor (GR) in fatty liver is still controversial (57, 70, 113). Surprisingly, we found that hepatic nuclear GR proteins were gene-dosage-dependently decreased by HNF4α deficiency in HFHS-fed mice. In parallel, HFHS-fed GR knockout mice had elevated hepatic levels of cholesterol and triglycerides and induction of key lipogenic genes. Peroxisome proliferator-activated receptor alpha (PPARα) and liver X receptor (LXR) are master regulators of lipid metabolism (88, 110). We found that GR and HNF4α cooperatively activated promoters of key lipid catabolic genes, whereas HNF4α and GR antagonized the induction of lipogenic genes by LXR and PPARα, respectively, suggesting that GR and HNF4α cooperatively protect against HFHS-induced fatty liver and hyperlipidemia.

## Results

### Differential effects of diets on blood lipid profiles in HNF4α heterozygote and knockout mice

Adult male HNF4α heterozygote and knockout mice fed different diets had distinct changes in blood lipid profiles 15 d after diet treatment. Under LFD, HNF4α heterozygote mice had similar blood lipids as wildtype (WT) mice, whereas HNF4α knockout mice had marked hypolipidemia, manifested by 72% lower triglycerides (TG) (Fig. 1A), 43% lower free fatty acids (Fig. 1B), and 44% lower total cholesterol (Fig. 1C) than WT mice, consistent with literature (47). Blood levels of high-density lipoprotein (HDL) cholesterol (Fig. 1D) and low-density lipoprotein (LDL)/very-low-density lipoprotein (VLDL) cholesterol (Fig. 1E) decreased similarly in the knockout mice, resulting in similar ratio of total/HDL cholesterol (Fig. 1F) in the LFD-fed WT and knockout mice.

**Figure 1.**
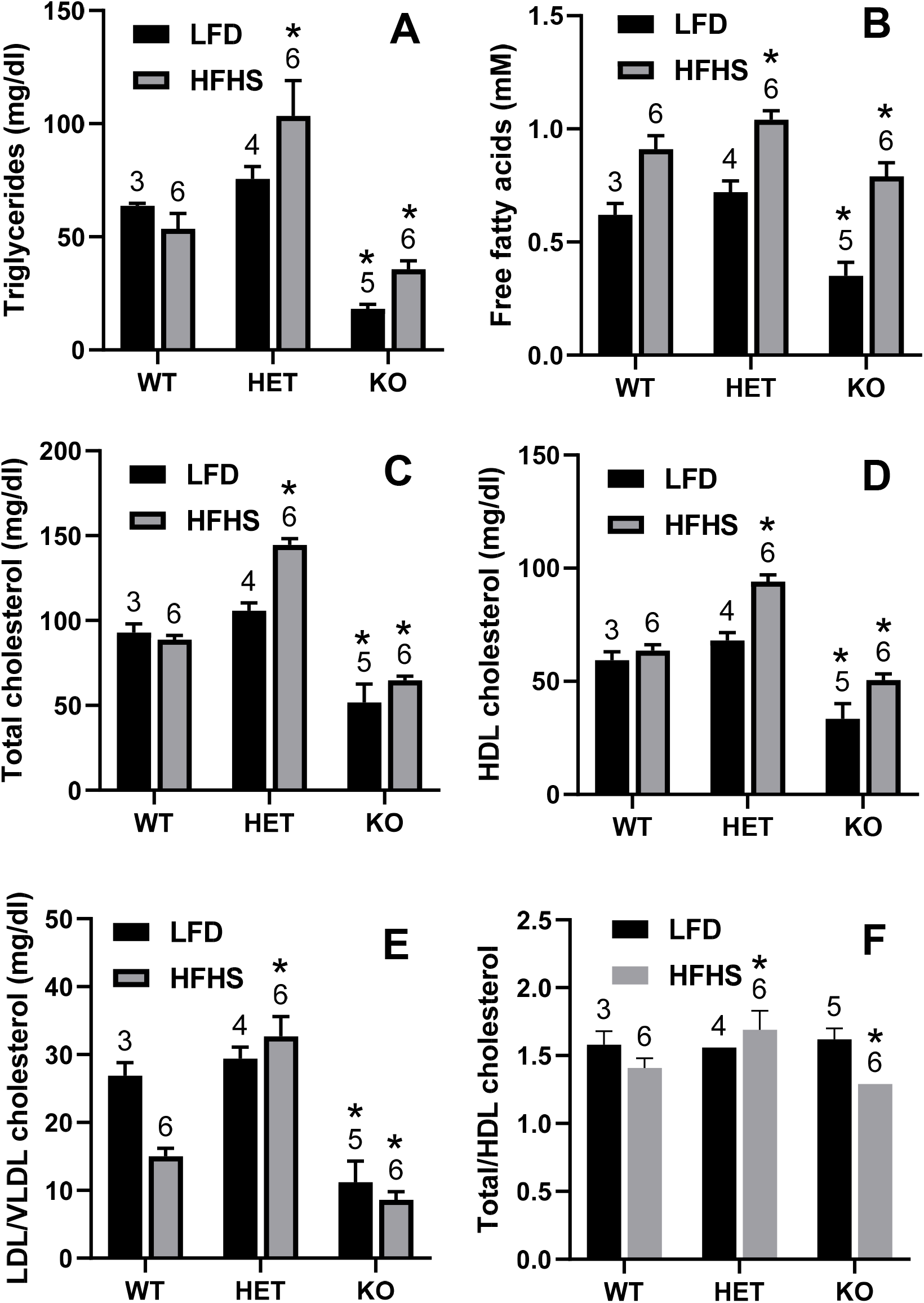
Blood levels of (A) triglycerides, (B) free fatty acids, (C) total cholesterol, (D) HDL cholesterol, (E) LDL/VLDL cholesterol, and (F) ratio of total/HDL cholesterol in adult male wildtype (WT), HNF4α heterozygote (HET), and HNF4α knockout (KO) mice. Mice were fed 15 d with low-fat diet (LFD, N=3-5 per group) or high-fat-high-sugar diet (HFHS) (N=6 per group). The number of animals in each group was labeled above the error bar. Mean ± SE. * p < 0.05 versus corresponding wildtype mice.

Under HFHS, HNF4α heterozygote mice had 93% higher TG (Fig. 1A), 14% higher free fatty acids (Fig. 1B), 63% higher total cholesterol (Fig. 1C), 48% higher HDL cholesterol (Fig. 1D), and 1.2 fold higher LDL/VLDL cholesterol (Fig. 1E) than WT mice, whereas HNF4α knockout mice still had mild hypolipidemia, namely 34% lower TG (Fig. 1A), 13% lower free fatty acids (Fig. 1B), 27% lower total cholesterol (Fig. 1C), 20% lower HDL cholesterol (Fig. 1D), and 42% lower LDL/VLDL cholesterol (Fig. 1E) than WT mice. Interestingly, the ratio of total/HDL cholesterol increased in the heterozygote but decreased in the knockout mice (Fig. 1F), suggesting that HFHS-fed heterozygotes have increased risk, whereas the knockouts have decreased risk of heart disease. Further studies were focused on understanding the distinct gene-dosage-dependent changes in lipid metabolism in the HFHS-fed HNF4α heterozygote and knockout mice.

### Changes in liver/body weight and blood levels of glucose and insulin in HFHS-fed mice

Three weeks after HFHS feeding, WT and HNF4α heterozygote mice tended to gain weight, whereas the knockout mice tended to lose weight (Table 1). The liver/body weight ratio was 18% and 47% higher in the heterozygote and knockout mice, respectively, compared to WT mice. Blood levels of glucose remained unchanged in heterozygote mice but decreased 35% in knockout mice. Consistent with changes in blood glucose, serum levels of insulin remained unchanged in heterozygote but decreased 36% in knockout mice. These data suggest enhanced hepatic insulin signaling in HFHS-fed HNF4α knockout mice. Interestingly, blood levels of ketone bodies, determined by β-hydroxybutyrate, were 63% and 29% higher in the heterozygote and knockout mice (Table 1).

**Table 1.** Changes of body weight, liver/body weight as well as blood levels of glucose, insulin, and ketone bodies in adult male wildtype, HNF4α heterozygote, and HNF4α knockout mice fed the high-fat-high-sugar diet for 15 d.

|  | Wildtype | Heterozygote | Knockout |
| --- | --- | --- | --- |
| Body weight (Day0) (g) | 23.6 ± 0.6 | 25.0 ± 0.5 | 23.7 ± 0.5 |
| Body weight (Day15) (g) | 24.9 ± 0.9 | 25.7 ± 0.5 | 21.7 ± 1.0 |
| Liver/body weight (g/100 g) | 4.25 ± 0.11 | 5.00 ± 0.19 * | 6.25 ± 0.21 * |
| Glucose (mg/dl) | 199 ± 5 | 180 ± 7 | 130 ± 9 * |
| Insulin (nM) | 0.21 ± 0.02 | 0.21 ± 0.06 | 0.13 ± 0.02 * |
| Ketone body (μM) | 205 ± 8 | 333 ± 46 * | 264 ± 11 * |

N=6 per group, mean ± SE. \* p < 0.05 versus wildtype mice.

### Differential effects of diets on hepatic lipid profiles in HNF4α heterozygote and knockout mice

Both HNF4α heterozygote and knockout mice fed the HFHS for 15 d had marked 139% and 118% increases in hepatic TG, respectively, compared to WT mice (Fig. 2A). Hepatic cholesterol was 30% higher in HFHS-fed knockout mice than WT mice (Fig. 2A). Compared to 15 d post-HFHS, hepatic TG increased markedly (135%) in WT males fed the HFHS for 6 weeks, and HNF4α heterozygote mice still had 46% higher TG than WT mice (Fig. 2B). Our data strongly suggest that partial loss of HNF4α after longer-term HFHS intake may be a key mechanism of the loss of the early resistance of the liver to HFHS-induced fatty liver.

**Figure 2.**
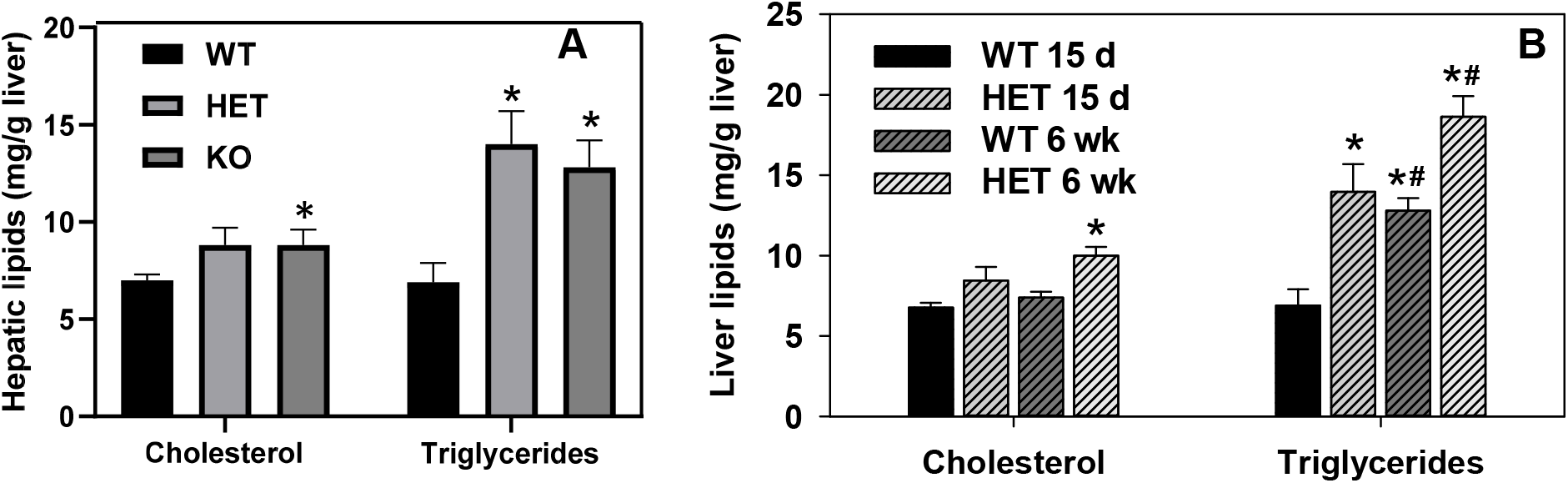
Hepatic lipids in adult male wildtype (WT), HNF4α heterozygote (HET), and/or HNF4α knockout (KO) mice. Mice were fed high-fat-high-sugar diet (HFHS) for (A) 15 d or (B) 6 weeks (6 wk). N=6 per group, mean ± SE. * p < 0.05 versus wildtype group. # p < 0.05 versus 15 d group.

### Changes in hepatic histology

Histology analysis (H&E staining) showed little changes of liver morphology in the HFHS-fed WT mice (Fig. 3B) compared to the LFD-fed WT mice (Fig. 3A). Compared to the corresponding WT mice, the HFHS-fed heterozygote mice (Fig. 3D) had more vacuolization of hepatocytes than the LFD-fed heterozygote mice (Fig. 3C). In contrast, the HFHS-fed knockout mice (Fig. 3F) had more marked enlargement and vacuolization of hepatocytes than the LFD-fed knockout mice (Fig. 3E). No obvious infiltration of inflammatory cells was observed in any of these livers.

**Figure 3.**
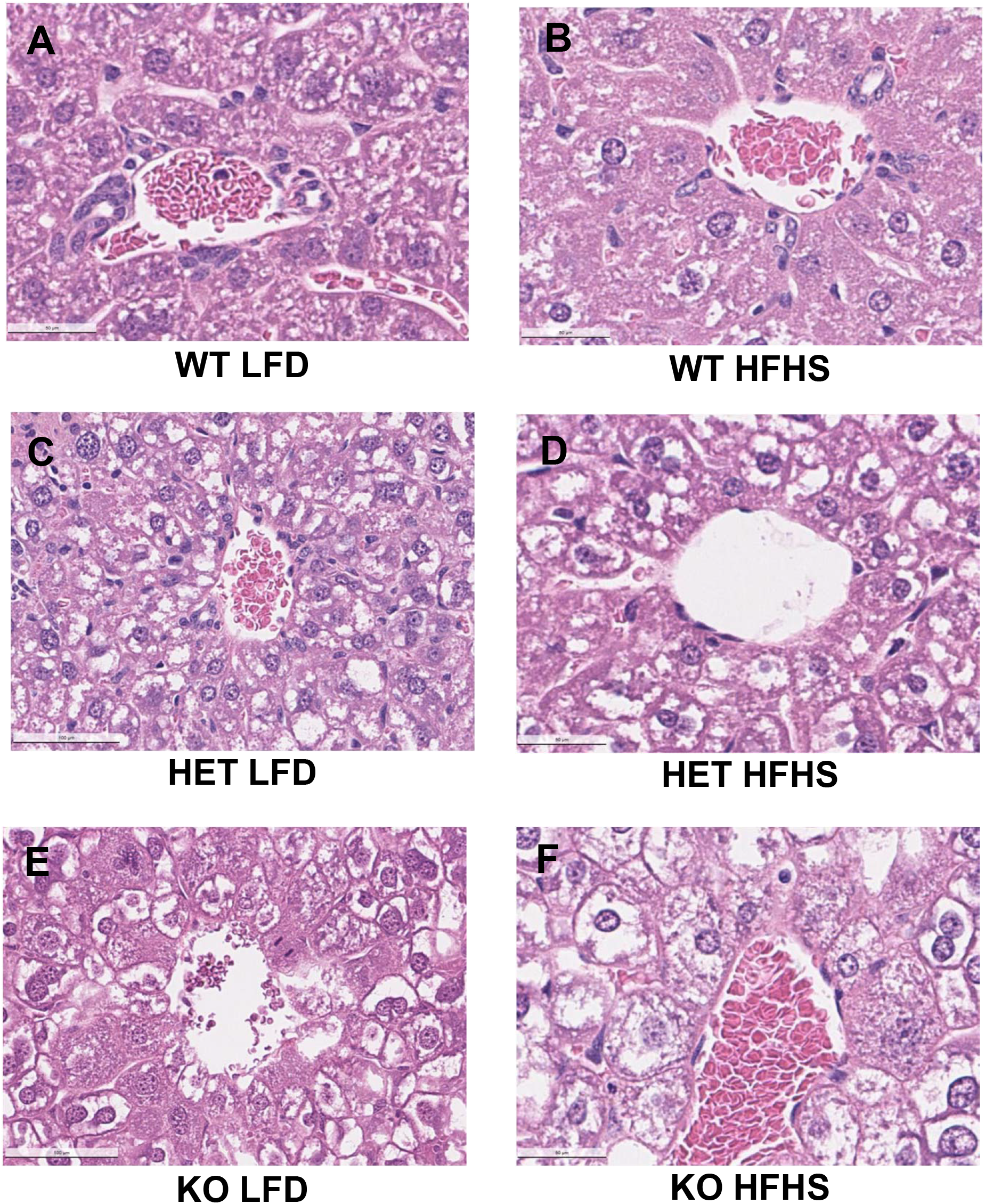
Liver histology in adult male wildtype (WT), HNF4α heterozygote (HET), and HNF4α knockout (KO) mice fed 15 d with low-fat diet (LFD) or high-fat-high-sugar diet (HFHS). H&E staining of paraffin embedded liver sections (5 μm and 400 × magnification). (A) WT LFD; (B) WT HFHS; (C) HET LFD; (D) HET HFHS; (E) KO LFD; and (F) KO HFHS.

### Changes in blood and hepatic levels of bile acids (BAs)

Biosynthesis of BAs from the cholesterol is a major pathway for the catabolism of cholesterol, whereas biliary excretion of BAs is essential for BA and cholesterol elimination and intestinal absorption of lipids. In the liver, the classical pathway of CYP7A1-CYP8B1 prefers the biosynthesis of hydrophilic cholic acid (CA), and the alternative pathway of CYP27A1-CYP7B1 prefer the hydrophobic chenodeoxycholic acid (CDCA). Both CYP27A1 and CYP7B1 are expressed in various tissues, whereas CYP7A1 and CYP8B1 are liver-specific (56). In rodents, the hydrophobic CDCA is largely converted to the highly hydrophilic muricholic acid (MCA) by CYP2C70, resulting in a much more hydrophilic BA pool than humans (128). Additionally, CDCA can be detoxified by CYP3A to more hydrophilic hyocholic acid (HCA) (17). The hydrophilic ursodeoxycholic acid (UDCA), a primary BA in the bear, is a minor form in mice. In the intestine, CA and CDCA are metabolized by bacteria to cytotoxic deoxycholic acid (DCA) and lithocholic acid (LCA), respectively. Most LCA can be efficiently sulfated and excreted from the intestine, whereas unconjugated DCA is reabsorbed in the large intestine and transported back to the liver, where it can be conjugated (71). In rodents, the taurine (T) conjugated DCA (T-DCA) can be converted back to T-CA via 7-hydroxylation by Cyp2a12 (50). Additionally, the bacteria-metabolized 7-ketoLCA can be absorbed and converted back to CDCA in the liver (38). Hyodeoxycholic acid (HDCA), generated by bacterial C-6 hydroxylation of LCA or dehydroxylation of HCA in small intestine, is a LXRα agonist (125). Approximately 95% of BAs, consisting of mainly CA, DCA, and CDCA in humans and CA and α/β-MCA in mice, are reabsorbed via enterohepatic circulation. Only ~5% BAs are synthesized *de novo* from cholesterol daily.

#### Changes of blood levels of BAs

compared to LFD-fed WT mice, HFHS-fed WT mice had much lower blood levels of total BAs (↓64%) (Fig. 4A), including the primary BAs CA (↓66%) MCA (↓75%), and UDCA (↓85%) and the secondary BAs 12-oxo-CDCA (↓92%) and HDCA (↓69%). HFD intake increases BA secretion and fecal levels of BAs in humans and rodents (135). Particularly, HFD selectively increases the concentration of T-CA in the bile (148). In contrast, HFD intake is associated with decreased blood levels of total and primary BAs in humans (135). Thus, the decreases of blood BAs in HFHS-fed WT mice is likely due to increased biliary secretion and fecal elimination of primary BAs.

**Figure 4.**
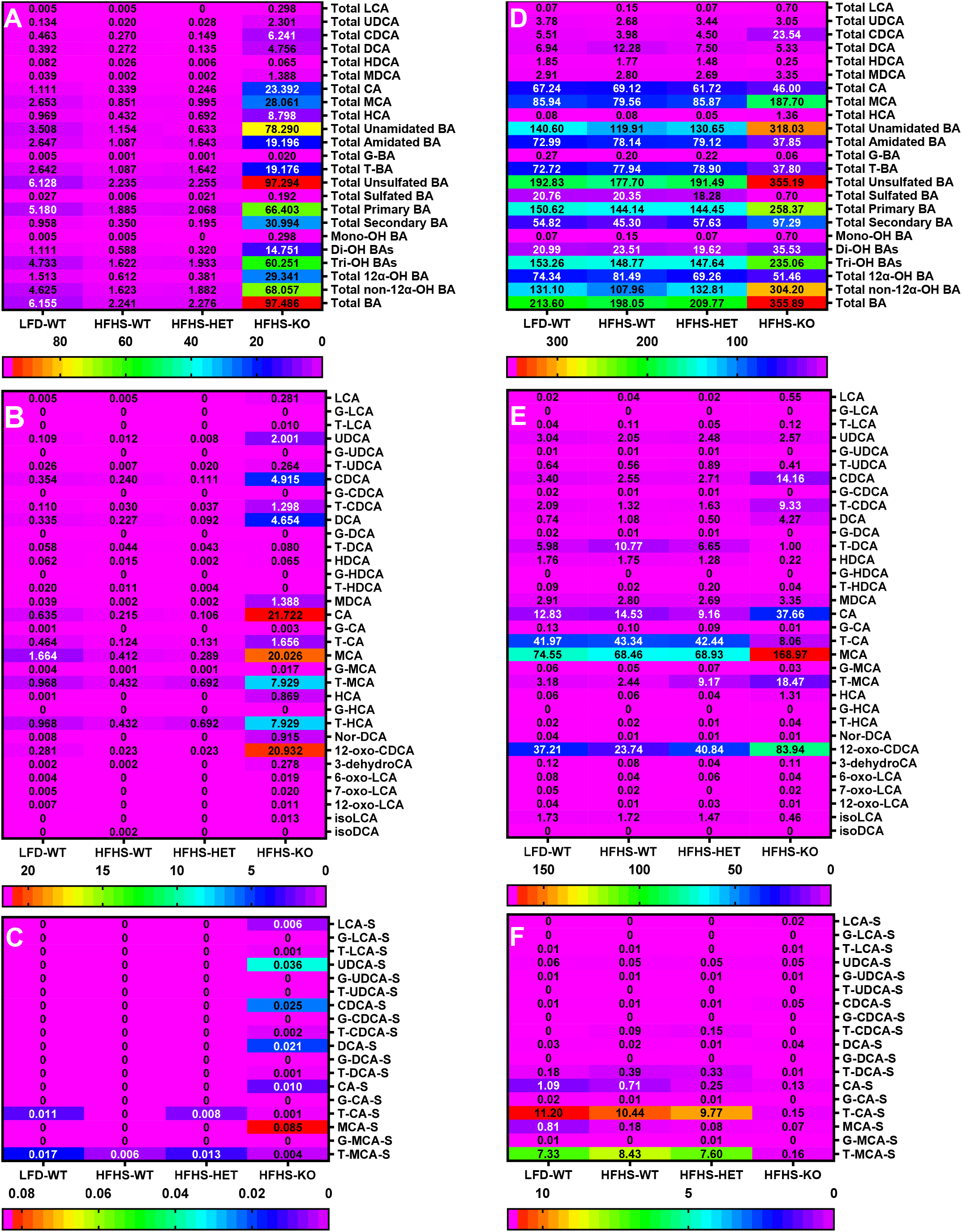
Serum and hepatic levels of bile acids in adult male wildtype (WT), HNF4α heterozygote (HET), and HNF4α knockout (KO) mice fed 15 d with low-fat diet (LFD) (N=3) or high-fat-high-sugar diet (HFHS) (N=6 per group). Bile acids were quantified by LC-MS/MS. (A) serum levels of total bile acids; (B) serum levels of non-sulfated bile acids; (C) serum levels of sulfated bile acids; (D) hepatic levels of total bile acids; (E) hepatic levels of non-sulfated bile acids; (F) hepatic levels of sulfated bile acids. The numbers in the heat maps represent mean bile acid concentrations (μM).

Interestingly, compared to HFHS-fed WT mice, the heterozygote mice had similar levels of total BAs, but 45%, 50%, and 54% less total unamidated BAs, total DCA, and CDCA, respectively (Fig. 4A). In contrast, the knockout mice had marked increases in total levels of unamidated BAs (↑77 fold), amidated BAs (↑18 fold), glycine-conjugated BAs (G-BAs) (↑37 fold), T-BAs (↑17 fold), unsulfated BAs (↑43 fold), sulfated BAs (↑30 fold), primary BAs (34 fold), secondary BAs (↑87 fold), mono-OH BAs (↑64 fold), Di-OH BAs (↑24 fold), Tri-OH BAs (↑36 fold), 12α-OH BAs (↑47 fold), and non-12α-OH BAs (↑40 fold), with a 42 fold increase in total BAs in the knockout mice (Fig. 4A). The knockout mice had dramatic increases in many primary and secondary BAs, namely CA (↑100 fold), T-CA (↑12 fold), MCA (↑48 fold), G-MCA (↑31 fold), T-MCA (↑17 fold), CDCA (↑19 fold), T-CDCA (↑42 fold), LCA (↑60 fold), UDCA (↑162 fold), T-UDCA (↑35 fold), DCA (↑19 fold), HDCA (↑3.5 fold), MDCA (↑887 fold), T-HCA (↑17 fold), 12-oxo-CDCA (↑903 fold), 3-dehydroCA (↑137 fold) (Fig. 4B). Additionally, certain sulfated (S) BAs, which were barely detectable in the WT mice, were significantly elevated in the knockout mice, such as UDCA-S (36 nM), CDCA-S (25 nM), DCA-S (21 nM), CA-S (10 nM), and MCA-S (85 nM) (Fig. 4C). Regarding the composition of total BAs, the HNF4α knockout mice had a marked 65%, 60%, and 58% decreases in % amidation of total BAs, % HCA, and % DCA respectively, whereas 1.1, 17, and 1.6 fold increases in % secondary BAs, % MDCA, and % UDCA, respectively. The overall hydrophobicity index (HI) of blood BAs remained unchanged in these mice.

#### Changes in hepatic bile acids in HFHS-fed heterozygote mice

Surprisingly, compared to the LFD-WT mice, HFHS-fed WT mice had little change in hepatic BAs (Fig. 4D-F), despite marked differences in blood BA profiles in these mice. Compared to HFHS-fed WT mice, most of the BAs remained unchanged in the heterozygote mice. Interestingly, the heterozygote mice had significant increases in T-MCA (↑2.8 fold) and T-HDCA (↑8.4 fold), without changes in hepatic total levels of MCA and HDCA (Fig. 4E). The percentage of total non-12α-hydroxylated BAs significantly increased in the heterozygote (63.1 ± 1.2) compared to WT (56.2 ± 2.1) mice, and the hydrophobicity index (HI) decreased in the heterozygote (0.01 ± 0.03) compared to WT (0.11 ± 0.02) mice. The lower hydrophobicity index in the heterozygote mice suggests a more hydrophilic BA pool, which may alter intestinal absorption of lipids (89).

#### Changes in hepatic bile acids in HFHS-fed knockout mice

compared to HFHS-fed WT mice, the knockout mice had a trend of decreased total CA (↓34%) but a marked 4.9 fold increase of total CDCA, which is associated with increases in free CDCA (↑4.5 fold), T-CDCA (↑6.0 fold), and oxo-CDCA (↑2.5 fold) (Fig. 4D, 4E). Hepatic T-MCA markedly increased 6.6 fold, and free MCA also tended to increase (p=0.055) in the knockout mice in which HNF4α was not knocked out until the adult age. The increases of MCA in the knockout mice is consistent with the marked cholestasis and the efficient enterohepatic recycling of MCA.

Regarding secondary BAs, HDCA markedly decreased by 86%, whereas HCA increased 15 fold in the knockout mice (Fig. 4E), suggesting decreased bacterial conversion of LCA to HDCA but increased CYP3A-catalyzed conversion of CDCA to HCA in the knockout mice. Total DCA tended to decrease in the knockout mice, with dramatic decrease of T-DCA (↓91%) and a trend of increase of free DCA. In contrast, hepatic levels of UDCA, LCA, and murideoxycholic acid (MDCA), a secondary bile acid produced from MCA, remained little changed.

Sulfation of BAs plays an important role in the detoxification of BAs. T-CA sulfate (T-CA-S) and T-MCA sulfate (T-MCA-S), two major sulfated BAs in WT mouse livers, were dramatically decreased 85 and 98%, respectively, in the knockout mice (Fig. 4F). Total sulfated BAs decreased 97%, whereas the unsulfated BAs doubled in livers of the knockout mice (Fig. 4D). Total amidated BAs tended to decrease (↓52%), whereas the total unamidated BAs increased 1.7 fold in the knockout mice. Additionally, hepatic total levels of G-BAs, a minor form of amidated BAs in rodents, were 71% lower in the knockout mice (Fig. 4D).

### Gene-dosage-distinct changes in hepatic transcriptome in HFHS-fed HNF4α heterozygote and knockout mice

#### A. Gene-dosage-distinct changes in genes essential for hepatocyte proliferation and differentiation (Fig. 5A)

We first conducted RNA-sequencing of pooled livers, followed by verification with real-time PCR using individual samples (N=4-6 per group) (Fig. 5). As expected, Hnf4a was gene-dosage-dependently decreased in the HNF4α heterozygote and knockout mice (Fig. 5A). HNF4α deficiency caused gene-dosage-dependent induction of HNF4g in HNF4α heterozygote (↑1.3 fold) and knockout (↑16.8 fold) mice, which might partially compensate for the loss of HNF4α. HNF4α is a well-established master regulator of hepatocyte differentiation and maturation, whereas HNF1β is critical for the differentiation of hepatoblasts into cholangiocytes (22). Our previous study found that Hnf1b was induced in HNF4α knockout mice (82). Interestingly, HNF1b was gene-dosage-dependently induced in HNF4α heterozygote (↑2 fold) and knockout (↑5.7 fold) mice. Hepatocytes and cholangiocytes have a common precursor of hepatoblasts. Cited2, a co-activator of HNF4α, plays a key role in liver development (107). Interestingly, cited2 was induced in heterozygote, but not knockout mice (Fig. 5A). The thyroid cancer-1 (C8orf4) inhibits self-renewal of liver cancer stem cells by suppressing NOTCH2 signaling (156). The NOTCH2-SOX9 signaling promotes biliary epithelial cell specification during bile duct development and cholangiocarcinogenesis (2, 132). Activation of Notch signaling in hepatocytes also promotes the secretion of osteopontin (SPP1), which promotes myofibroblast differentiation and liver fibrosis (2). We found 71% down-regulation of C8orf4 in the knockout mice and a clear trend of gene-dosage-dependent induction of Sox9, Spp1, and transforming growth factor beta (Tgfb) in HNF4α heterozygote (↑94%, 68%, and 25%, respectively) and knockout (↑6.1, 9.8, and 1.9 fold, respectively) mice (Fig. 5A).

**Figure 5.**
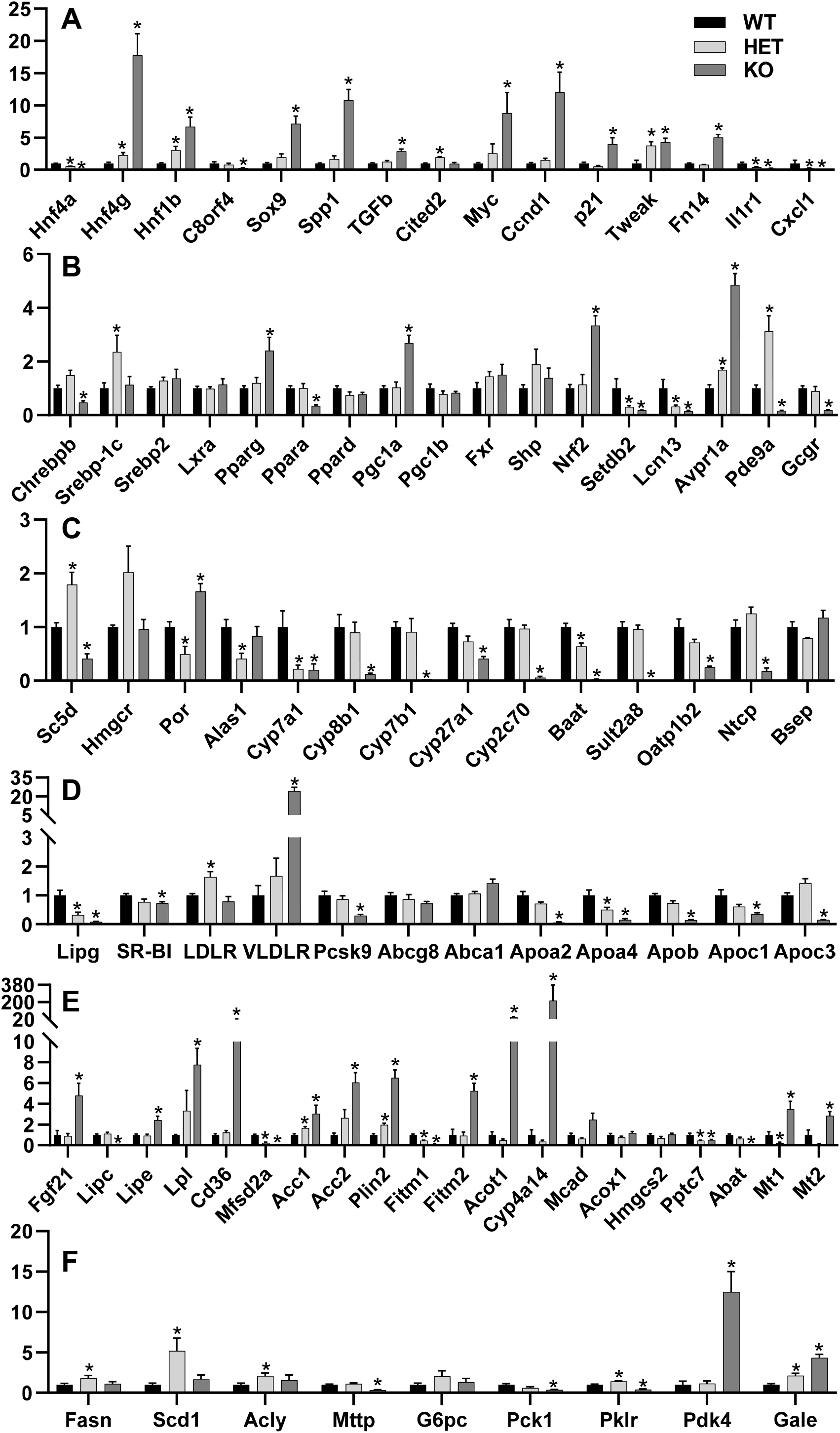
Real-time PCR quantification of hepatic mRNAs in adult male wildtype, HNF4α heterozygote (heterozygote), and HNF4α knockout (knockout) mice fed the high-fat-high-sugar diet (HFHS) for 15 d. (A) genes important in the differentiation and proliferation of hepatocytes and cholangiocytes; (B) transcriptional regulators; (C) genes important for cholesterol and bile acid metabolism; (D) genes important for apolipoprotein metabolism; (E) genes important in lipid metabolism; and (F) genes important for the metabolism of fatty acids and sugar. N=4-6, mean ± SE. Data were normalized to peroxiredoxin 1 (Prxd1), with wildtype values set at 1.0. * p < 0.05 versus wildtype control mice.

Total loss of HNF4α causes marked hepatocyte proliferation (10). We found gene-dosage-dependent induction of c-Myc and cyclin d1 (Ccnd1) in HNF4α heterozygote and knockout mice (Fig. 5A). Induction of TNF-like weak inducer of apoptosis (TWEAK) ligand and its receptor fibroblast growth factor-inducible 14 (Fn14) plays a key role in hepatocyte proliferation during chronic liver diseases (26). Interestingly, the tweak ligand was strongly induced in both the heterozygote (↑2.8 fold) and knockout (↑3.3 fold) mice, whereas the Fn14 receptor, which is not expressed in normal hepatocytes, was only induced in the knockout mice (↑4.0 fold). Additionally, the CDK inhibitor p21, an HNF4α-target gene (53), tended to be down-regulated in the heterozygote mice but strongly induced 3.0 fold in the knockout mice (Fig. 5A). Taken together, these data clearly demonstrate a critical gene-dosage-dependent role of HNF4α in maintaining hepatocyte differentiation and suppressing hepatocyte proliferation, cholangiocyte transdifferentiation, and liver fibrosis during HFHS intake.

Acute loss of HNF4α promotes rapid hepatocyte proliferation without the induction of inflammatory cytokines (10); the underlying mechanism remains unknown. Chemokine (C-X-C motif) ligand 1 (CXCL1) and interleukin-1 receptor 1 (IL1R1) play key roles in regulating hepatic proinflammatory responses (14, 40). We found gene-dosage-dependent critical roles of HNF4α in regulating expression of Cxcl1 and Il1r1 in mouse livers: Cxcl1 was 79% and 95% lower, whereas Il1r1 was 60% and 81% lower in the heterozygote and knockout mice (Fig. 5A). Hepatic down-regulation of Cxcl1 and Il1r1 may play important roles in preventing the infiltration of inflammatory cells and activation of the inflammatory kinase pathways in the HNF4α knockout mice (10) (Fig. 3).

#### B. Gene-dosage-distinct changes in transcriptional regulators (Fig. 5B)

Carbohydrate-responsive element-binding protein (ChREBP), SREBP-1, and LXRs are key lipogenic transcription factors. HNF4α is essential for transactivating ChREBPβ, a constitutive-active isoform of ChREBP (85). HNF4α knockout mice had 53% lower Chrebpβ than WT mice (Fig. 5B). In contrast, HNF4α heterozygote mice had 1.4 fold higher Srebp-1c and a trend of higher (48%) Chrebpβ than WT mice. Peroxisome proliferator-activated receptor (Ppar) family members regulate genes in lipid and carbohydrate metabolism. Hepatic mRNA expression of Srebp2, Lxrα, and Ppard remained unchanged in these mice (Fig. 5B). In contrast, Ppara was down-regulated 65%, whereas Pparg was up-regulated 1.4 fold in the knockout mice. PPARγ coactivator 1-alpha (PGC1α) is a master regulator of mitochondrial biogenesis and gluconeogenesis. PGC1a, but not PGC1b, was induced 1.7 fold in the knockout mice. The BA receptor farnesoid X receptor (FXR) and its target gene orphan nuclear receptor small heterodimer partner (SHP) play key roles in BA and lipid metabolism. SHP is induced in NAFLD, and SHP induction promotes steatosis but inhibits inflammation, partly via induction of PPARγ and suppression of NF-kB (83). FXR and SHP tended to be higher in the heterozygote and knockout mice. Consistent with previous report of induction of antioxidative genes (82), NF-E2-related factor-2 (Nrf2), a master regulator of antioxidative responses, was induced 2.3 fold in the knockout mice (Fig. 5B). Induction of the epigenetic modifier SetDB2 by GR ameliorates fatty liver (113). Lipocalin 13 (LCN13) protects against fatty liver by inhibiting lipogenesis and stimulating fatty acid (FA) β-oxidation (FAO) (121). Plasma vasopressin (VP) is increased in diabetic patients and promotes fatty liver by activating hepatic arginine VP receptor 1A (Avpr1a) (49, 84). Hepatic expression of Setdb2, Lcn13, and Avpr1a were highly gene-dosage-dependent on HNF4α (Fig. 5B): Setdb2 was 69% and 82% lower, Lcn13 was 69% and 86% lower, whereas Avpr1a was 69% and 3.8 fold higher, in the heterozygote and knockout mice than WT mice, respectively. Loss of hepatic glucagon receptor (GCGR) lowers blood glucose and increases insulin sensitivity (78). Inhibition of the cGMP-specific phosphodiesterase 9A (PDE9A) decreases gluconeogenesis and blood glucose levels (119). Hepatic Gcgr and Pde9a were markedly down-regulated in knockout mice but unchanged or up-regulated in the heterozygote mice (Fig. 5B), which may contribute to the decreased blood glucose levels in the knockout mice (Table 1).

#### C. Gene-dosage-dependent changes of genes important in cholesterol and BA metabolism (Fig. 5C & 5D)

Lathosterol 5-desaturase (Sc5d), essential for cholesterol synthesis (66), was 79% higher in heterozygote but 59% lower in knockout mice than WT mice (Fig. 5C). HMG-CoA reductase (HMGCR), the rate-limiting enzyme for cholesterol biosynthesis, was induced 1.0 fold in the heterozygote but remained unchanged in the knockout mice. Cytochrome P450 reductase (POR) is essential in lipid metabolism by maintaining the activities of all P450s (92), whereas the liver-predominant aminolevulinic acid synthase 1 (ALAS1) is rate-limiting in heme synthesis whose deficiency contributes to mitochondrial dysfunction and NAFLD progression. Interestingly, Por and Alas1 mRNAs were 51% and 59% lower, respectively, in heterozygote mice than WT mice (Fig. 5C). Importantly, Por heterozygote mice have ~50% decrease in Por activity (44), suggesting decreased POR activity in heterozygote mice.

CYP7A1 is important in cholesterol catabolism and prevention of HFD-induced fatty liver (74). Cyp7a1 was down-regulated in both heterozygote (↓78%) and knockout (↓80%) mice, whereas only the knockout mice had significant down-regulation of Cyp8b1 (↓91%), Cyp7b1 (↓97%), Cyp27a1 (↓59%), and Cyp2c70 (↓94%) (Fig. 5C). Consistent with previous report (54), bile acid-CoA:amino acid N-acyltransferase (BAAT), the key enzyme for amino acid conjugation of bile acids, was down-regulated 98% in the knockout mice. Sulfotransferase 2a8 (Sult2a8), a newly identified major hepatic BA sulfonating enzyme in mice (31), was dramatically down-regulated (↓99%) in the knockout mice. Consistent with a previous report (82), expression of Na+-taurocholate cotransporting polypeptide (NTCP) and organic anion transporting polypeptide 1b2 (Oatp1b2), key uptake transporters for conjugated and unconjugated BAs (23), were markedly down-regulated 82% and 75%, respectively, in the knockout mice. In contrast, expression of bile salt export pump (BSEP) remained unchanged in the knockout mice (Fig. 5C).

Hepatic endothelial lipase (EL/LIPG) reduces blood levels of HDL cholesterol by promoting the HDL catabolism and hepatic uptake of HDL cholesterol (129). Lipg was gene-dosage-dependently down-regulated in the heterozygote (↓68%) and knockout (↓92%) mice (Fig. 5D). Scavenger receptor class B, type I (SR-BI), LDL receptor (LDLR), and VLDL receptor (VLDLR) are essential for hepatic uptake of HDL, LDL, and VLDL cholesterol, respectively. Ldlr was 64% higher, and Vldlr also tended to be higher in the heterozygote mice. In contrast, SR-BI was slightly decreased 27%, whereas Vldlr, a PPARα/PPARγ-target gene (130), was remarkably induced 23 fold in the knockout mice. Although Ldlr mRNA was not significantly altered in the knockout mice, proprotein convertase subtilisin/kexin type 9 (PCSK9), which plays a key role in the degradation of LDLR protein, was markedly decreased 70%, suggesting that hepatic protein levels of LDLR might be elevated in the knockout mice. Thus, the heterozygote and knockout mice likely have decreased hepatic uptake of HDL but increased uptake of VLDL/LDL cholesterol. Additionally, hepatic expression of the biliary cholesterol efflux transporter Abcg8 remained unchanged, whereas the basolateral cholesterol efflux transporter Abca1 tended to be higher in the knockout mice (Fig. 5D). The apolipoproteins Apo-AI and Apo-AII have major roles in regulating HDL cholesterol and lipid metabolism (104). Chow-fed Apoa2-null mice have 64% and 32% decreases in blood levels of HDL cholesterol and free fatty acids, without changes in non-HDL cholesterol and triglycerides (141). Consistent with the previous report of chow-fed HNF4α knockout mice (47), HFHS-fed HNF4α knockout mice had dramatic down-regulation of Apoa2 (↓94%), Apoa4 (↓85%), Apob (↓86%), Apoc1 (↓65%), and Apoc3 (↓85%) (Fig. 5D), but little changes in Apoa1 and Apoe (data not shown).

#### D. Gene-dosage-dependent changes of genes important in lipid metabolism (Fig. 5E)

Fibroblast growth factor 21 (FGF21) is a hepatokine that increases insulin sensitivity and regulates lipolysis in white adipose tissue (51). Hepatic Fgf21 was 3.8 fold higher in the knockout mice (Fig. 5E). The liver-predominant hepatic lipase (HL/LIPC) and fat-predominant lipoprotein lipase (LPL) mediate lipoprotein hydrolysis to provide free fatty acids for energy and storage. Loss of Lipc protects against, whereas liver-specific overexpression of LPL promotes diet-induced obesity and hepatic steatosis (19, 60). In contrast, hepatic overexpression of the hormone-sensitive lipase (HSL/LIPE) ameliorates fatty liver by promoting lipolysis and fatty acid oxidation (109). Interestingly, Lipc and Lipe were 94% lower and 1.4 fold higher in the knockout mice, which may help ameliorate fatty liver in these mice. In contrast, Lpl tended to be gene-dosage-dependently increased in the heterozygote (↑2.3 fold) and knockout (↑6.8 fold) mice (Fig. 5E). The free fatty acids released by these lipases are taken up into liver by fatty acid translocase (FAT/CD36). Hepatocyte-specific disruption of CD36 attenuates fatty liver and improves insulin sensitivity in HFD-fed mice, with large decreases in hepatic oleic acid without change in the essential ω3 FAs (142). In contrast, major facilitator super family domain containing 2a (MFSD2A) is a key transporter for ω3 FAs which is essential for prevention of fatty liver (95). Cd36 was strongly induced 20 fold in the knockout mice, whereas Mfsd2a was 77% and 94% lower in the heterozygote and knockout mice than WT mice, respectively, which may result in marked increase in hepatic uptake of oleic acid in the knockout mice but decreases in the uptake of ω3 FAs in the heterozygote and knockout mice.

The FAs taken up into liver will be utilized or stored. We found that the heterozygote and knockout mice had gene-dosage-dependent induction of key genes that inhibit lipid catabolism (Fig. 5E). Acetyl-CoA carboxylase (ACC) Acc1 and Acc2 are key lipogenic genes by inhibiting β-oxidation and promoting *de novo* lipogenesis. Perilipin 2 (PLIN2), a lipid droplet protein highly up-regulated in steatotic livers (13), promotes fatty liver and fibrosis (94). HNF4α had a gene-dosage-dependent critical role in preventing hepatic up-regulation of Acc1, Acc2, and Plin2 (Fig. 5E): Acc1 was 65% and 2.0 fold higher, Acc2 was 1.6 and 5.0 fold higher, whereas Plin2 was 98% and 5.5 fold higher, in HFHS-fed HNF4α heterozygote and knockout mice than WT mice, respectively. The muscle-predominant fat storage-inducing transmembrane protein 1 (FITM1) promotes the formation of smaller lipid droplets, likely to turn over FAs for mitochondrial β-oxidation (43). In contrast, overexpression of FITM2 increases lipid droplets and triglycerides in mouse liver (58). Interestingly, Fitm1 was 56% and 90% lower in heterozygote and knockout mice than WT mice, respectively, whereas Fitm2 was 4.2 fold higher in the knockout mice. FAs must be converted to acyl-CoAs for mitochondrial FAO. Acyl-CoA thioesterase 1 (ACOT1) hydrolyzes acyl-CoAs back to CoA and free FAs which promote fatty liver but inhibit inflammation and oxidative stress by activating PPARα (35). Additionally, the PPARα-target genes, namely the microsomal Cyp4a14, the peroxisomal acyl-coenzyme A oxidase 1 (ACOX1), as well as the mitochondrial enzymes medium-chain acyl-CoA dehydrogenase (MCAD), and hydroxymethylglutaryl-CoA synthase, mitochondrial (HMGCS2) play key roles in FAO (88). All these PPARα-target genes tended to be lower in the heterozygote mice, whereas Cyp4a14 was markedly induced (↑213 fold) and MCAD tended to be induced (↑1.5 fold, p = 0.056) in the knockout mice (Fig. 5E). The trend of induction of MCAD may be due to induction of PGC1a (Fig. 5B) which can strongly induce MCAD independent of HNF4α (111). Pptc7 is a newly identified essential phosphatase for promoting mitochondrial metabolism and biogenesis (96). Interestingly, Pptc7 was down-regulated in both the heterozygote (↓56%) and knockout (↓49%) mice (Fig. 5E). The gut-derived γ-Aminobutyric Acid (GABA) is accumulated in liver disease and contributes to hepatic encephalopathy in patients with cirrhosis (97). 4-aminobutyrate aminotransferase (ABAT), a key enzyme for GABA catabolism and mitochondrial nucleoside metabolism (9), tended to be gene-dosage-dependently decreased in the heterozygote and knockout mice (↓94%). Interestingly, GABA potently protects against severe liver injury (46). Thus, the dramatic down-regulation of ABAT in the knockout mice may impair mitochondrial nucleoside metabolism but protect the knockout mice from liver injury via elevated GABA. Metallothionein protects against HFD-induced fatty liver (114). Metallothionein-1 (Mt1) and Mt2 are the most markedly down-regulated genes in the HFD-fed obesity-prone C57/BL6 mice (138). Mt1 and Mt2 were markedly decreased 79% and 91%, respectively, in heterozygote mice but increased 2.5- and 1.9-fold in knockout mice (Fig. 5E), which may be due to stress responses such as activation of Nrf2 and FXR by cholestatic liver injury in these mice (82, 134).

#### E. Differential alterations of key genes for lipogenesis and sugar metabolism (Fig. 5F)

HNF4α heterozygote mice had induction of key lipogenic enzymes FA synthase (Fasn) (↑79%) and stearoyl-CoA desaturase-1 (Scd1) (↑4.2 fold). ATP citrate lyase (ACLY), a Srebp-1-target key enzyme in *de novo* FA and cholesterol synthesis (21), was 1.1 fold and 56% higher in the heterozygote and knockout mice. Additionally, knockout mice had 68% down-regulation of microsomal triglyceride transfer protein (Mttp) (Fig. 5F).

The metabolism of sugar and lipids are closely inter-related. Liver-type pyruvate kinase (PKLR), essential in glycolysis and lipogenesis (16), was 40% higher in heterozygote but 63% lower in knockout mice (Fig. 5F). Hepatic Pyruvate dehydrogenase kinase 4 (PDK4) levels are highly induced in human patients with NASH, whereas deletion of Pdk4 improves fatty liver in mice (151). Pdk4 was markedly induced 11.5 fold in the knockout mice. Interestingly, HNF4α deficiency in HFHS-fed mice had distinct effects on phosphoenolpyruvate carboxykinase (PEPCK/PCK1) and glucose-6-phosphatase catalytic subunit (G6PC), two key enzymes for gluconeogenesis. Pepck tended to be gene-dosage-dependently down-regulated (↓65% in knockout mice), whereas G6pc tended to be up-regulated in heterozygote and knockout mice. Pepck promotes gluconeogenesis but protects against fatty liver (120). The marked down-regulation of Pepck likely contributes to decreased blood glucose and accumulation of lipids in the HFHS-fed knockout mice. Hepatic expression of UDP-galactose-4-epimerase (GALE), the last enzyme in the conversion of UDP-galactose to UDP-glucose, is induced after long-term HFHS feeding (157). Hepatic induction of GALE impairs insulin sensitivity (157). Interestingly, Gale was gene-dosage-dependently induced in the heterozygote (↑1.1 fold) and knockout (↑ 3.3 fold) mice (Fig. 5F), suggesting a key role of HNF4α in preventing hepatic induction of GALE during HFHS intake.

### Western blot quantification of protein expression of genes essential for lipid metabolism in HFHS-fed HNF4α heterozygote and knockout mice

HNF4α heterozygote and knockout mice had expected partial and complete loss of Hnf4α proteins (Fig. 6). Consistent with literature (10), nuclear β-catenin was markedly increased in knockout mice. Activation of β-catenin increases insulin sensitivity and strongly inhibits Srebp-1c expression (1, 150), which may explain a lack in the induction of Srebp-1c mRNA in HNF4α knockout mice (Fig. 6A). The Srebp-1 protein is synthesized as a precursor form (110 kDa) anchored in the endoplasmic reticulum and nuclear membranes. After proteolytic cleavage, the mature active form (50 kDa) of Srebp-1c migrates into the nucleus (11). Nuclear precursor form (110 KD) of Srebp-1c remained unchanged in HNF4α heterozygote mice but decreased in HNF4α knockout mice. In contrast, nuclear mature form of Srebp-1c (50 KD) markedly increased in HNF4α heterozygote mice. AMPK is a key regulator of lipid metabolism, partly via phosphorylating and inhibiting ACC (79). Hepatic AMPK activity is decreased in obesity and diabetes. We found that nuclear phosphorylated AMPK (p-AMPK) was unchanged in HNF4α heterozygote, but decreased markedly in HNF4α knockout mice (Fig. 6). Consistent with the gene-dosage-dependent induction of Acc mRNAs (Fig. 5E), cytosolic Acc proteins increased moderately in HNF4α heterozygote and dramatically in HNF4α knockout mice (Fig. 6). Parallel with changes in p-AMPK, phosphorylated Acc (p-Acc) was unchanged in heterozygote but decreased in HNF4α knockout mice. Surprisingly, Fasn proteins increased markedly in both HNF4α heterozygote and knockout mice (Fig. 6), despite a lack in the induction of Fasn mRNA in HNF4α knockout mice (Fig. 5F). Activation of AMPK depletes Fasn protein in the cytosol (98). Thus, the marked decrease in p-AMPK may be responsible for increased cytosolic Fasn protein in HNF4α knockout mice. 11β-hydroxysteroid dehydrogenase type 1 (HSD11B1) is responsible for local activation of GR (15). Hepatic Gr and Hsd11b1 mRNAs were unchanged in HNF4α heterozygote and knockout mice (data not shown). Surprisingly, nuclear GR proteins were moderately and dramatically decreased in HNF4α heterozygote and knockout mice, whereas cytosolic GR proteins were unchanged in these mice (Fig. 6).

**Figure 6.**
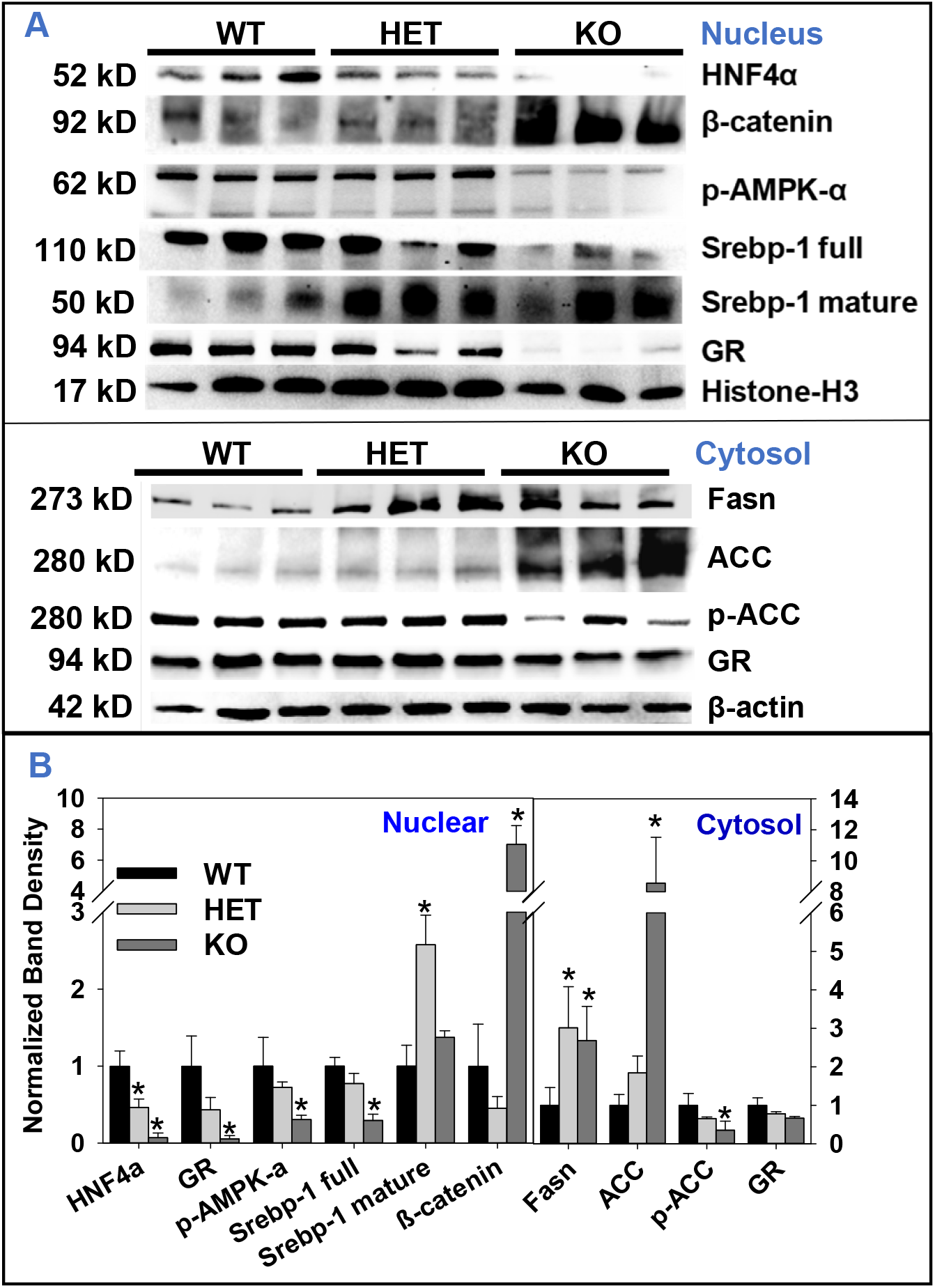
Western blot quantification of proteins in hepatic nuclear (top) and cytosolic (bottom) extracts in adult male wildtype (WT), HNF4α heterozygote (HET), and HNF4α knockout (KO) mice fed the high-fat-high-sugar diet for 15 d. (A) Gel image. (B) Band density normalized to histone H3 (nucleus) or β–actin (cytosol). N=3 per group, Mean ± SE. * p < 0.05 versus wildtype mice.

### Suppression of mouse and human SREBP-1C promoter by HNF4α

SREBP-1C and LXR are two master regulators of lipogenesis. The basal expression of SREBP-1C is dependent on LXR, a key lipogenic nuclear receptor that is activated by oxysterol (110). Without a DNA-binding domain, the orphan receptor SHP mainly acts as a co-repressor (152). Bile acids lower hepatic TG via FXR-SHP-LXR-SREBP-1c pathway (140). We found that HNF4α gene-dosage-dependently suppressed the LXR-mediated transactivation of reporters for mouse and human SREBP-1C promoter in HEK293 cells (Fig. 7A). An E363K mutation in the AF-2 activation domain of HNF4α does not affect its cellular protein expression (55), but blocks its interaction with SHP (69). In hepatoma cells, SHP potently inhibits the transactivation of Srebp-1c by LXRα (110). In contrast, we found that SHP only weakly inhibited LXRα-transactivation of Srebp-1c promoter in HEK293 cells that lack HNF4α expression (Fig. 7B). Interestingly, WT HNF4α, but not the SHP-interaction-defective E363K mutant HNF4α, cooperated with SHP to synergistically inhibit the LXRα-transactivation of human SREBP-1C (Fig. 7B) and the basal mouse Srebp-1c promoter activities (Fig. 7C). Thus, SHP mediates the strong inhibition of LXR-transactivation of human SREBP-1C and mouse Srebp-1c by HNF4α.

**Figure 7.**
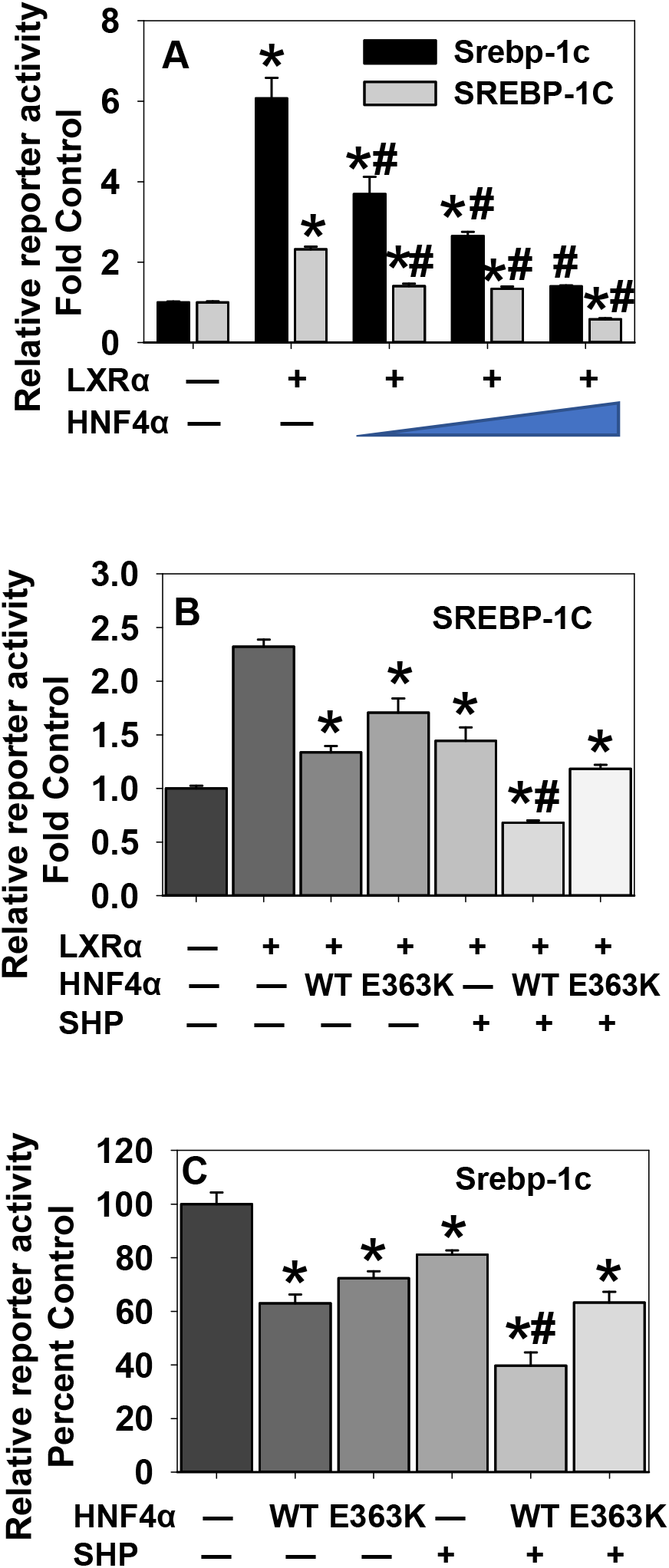
Dual-luciferase assays of regulation of mouse Srebp-1c and human SREBP-1C promoters. Assays were conducted 24 h after HEK293 cells (A & B) or HepG2 cells (C) were transfected with firefly reporter vectors, pRL-CMV, and expression vectors for HNF4α, LXRα, or SHP. N=4, mean ± SE. * p < 0.05 versus control; # p <0.05 versus (A) LXR, (B & C) HNF4α or SHP alone.

### Crosstalk of GR with HNF4α and PPARα in regulation of hepatic gene expression

GR and HNF4α can cooperate to induce CYP2A6 and PEPCK in humans (100, 126). Additionally, GR can gene-specifically enhance or antagonize the transactivation by PPARα (102, 108), a master regulator of lipid metabolism. Thus, we hypothesized that down-regulation of GR signaling (Fig. 6) plays a key role in the dysregulation of HNF4α- and PPARα-target genes in HNF4α heterozygote and knockout mice. Histone H3 trimethylation at lysine-4 (H3K4me3) is a marker for active transcription, whereas H3K4me1 is a marker for active enhancer (61). The public datasets of ChIP-sequencing of H3K4me3 (GSM769014), H3K4me1 (GSM722760), HNF4α (GSM1390711), GR in fed state (GR_Fed, GSM1936962), and GR in 24h-fasted state (GR_Fast, GSM1936964) in WT mouse livers were uploaded into the Integrative Genomics Viewer (IGV) software (112) to visualize their DNA-binding in each gene locus. We found both HNF4α peaks and enhanced GR-binding after fasting in promoters of Lcn13, Cyp7a1, Mt1, and Apoc3, intron5 of Setdb2, intron1 of Por, and intron9 of Alas1, except that only HNF4α was found in Fitm1 exon1 and GR was found in Mfsd2a intron2 (Fig. 8A).

**Figure 8.**
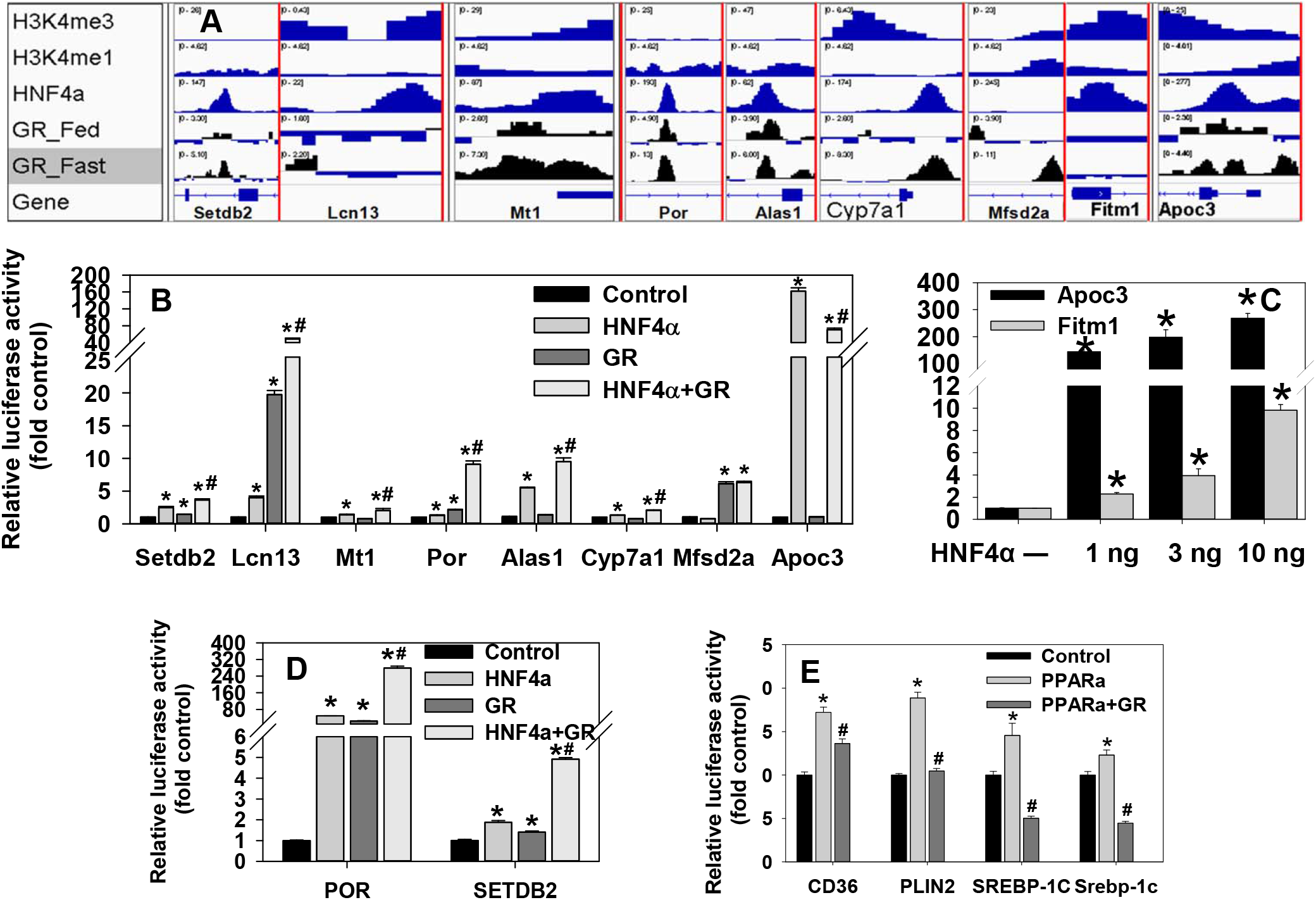
Interactions of GR with HNF4α and PPARα in gene regulation. (A) Analysis of ChIP-sequencing data of DNA-binding of epigenetic signatures, HNF4α, and GR in gene loci in wildtype mouse liver. (B - E) Dual-luciferase assays of promoter/intron activities of mouse genes (B-C), human POR and SETDB2 (D), as well as human CD36, human PLIN2, human SREBP-1C, and mouse Srebp-1c (E) genes in HEK293 cells 24 h after transfection with firefly reporter vectors, pRL-CMV, and expression vectors for HNF4α, GR (with 10 nM dexamethasone), or PPARα (with 10 μM clofibrate). N=4, mean ± SE. * p < 0.05 versus control; # p <0.05 versus HNF4α or PPARα alone.

We generated reporter vectors by PCR-cloning the DNA elements in genes that show strong binding of GR and/or HNF4α in the ChIP-sequencing data (Fig. 8A), into the pGL3-Basic vector. We conducted dual-luciferase assays 24 h after the HEK293 cells were co-transfected with the reporter vectors and expression vectors of HNF4α and/or GR (with 10 nM dexamethasone). HNF4α and GR synergistically/additively transactivated Setdb2, Lcn13, Mt1, Por, Alas1, and Cyp7a1 (Fig. 8B). In contrast, only GR transactivated Mfsd2a, whereas GR antagonized the strong transactivation of Apoc3 by HNF4α (Fig. 8B). Moreover, HNF4α and GR also synergistically transactivated human POR and SETDB2 introns (Fig. 8D). Additionally, HNF4α highly gene-dosage-dependently transactivated Fitm1 promoter, whereas the Apoc3 promoter was very strongly transactivated by merely 1 ng of HNF4α expression vector (Fig. 8C). Thus, the dramatic down-regulation of Apoc3 in HNF4α knockout mice (Fig. 6D) may be due to the loss of HNF4α as an obligatory transactivator, whereas the trend of induction of Apoc3 in heterozygote mice might be due to the decreased suppressing effects of GR on Apoc3.

Previous study demonstrates that certain PPARα-target genes are highly induced in chow-fed HNF4α knockout mice despite hepatic down-regulation of PPARα (47). PPARα induces the key lipogenic transcription factor Srebp-1c by directly binding to its promoter (32). We found that GR completely blocked the transactivation of human SREBP-1C and mouse Srebp-1c promoters by PPARα (Fig. 8E). Activation of PPARα induces key lipogenic genes PLIN2 and CD36 in mouse and human hepatocytes (27, 115, 131). We found that GR completely and partially blocked, respectively, the activation of intron1 of human PLIN2 and CD36 by PPARα (Fig. 8E).

### Expression of hepatic GR- and PPARα-target genes in WT mice fed HFHS for 15 d

Compared to LFD-fed mice, despite a 1.2 fold induction of Srebp-1c, Scd1 was markedly down-regulated 89%, and PLIN2 was only moderately induced 1.2 fold in the HFHS-fed WT mice (Supplemental Fig. 1) (https://figshare.com/s/4a15e7089b4744443033). Hmgcr and Cyp7a1, key enzymes for the synthesis and degradation of cholesterol, was 4.7 and 4.1 fold higher, and Alas1 and Por were 13.3 and 6.6 fold higher, suggesting that the turnover of cholesterol may be increased by short-term HFHS intake. High-fructose diet feeding for 2-3 weeks markedly suppresses hepatic PPARα signaling in rats and mice (59, 127). Consistently, hepatic expression of PPARα and the marker genes for PPARα activation, namely Cyp4a14 and acyl-CoA oxidase 1 (Acox1) were not induced 15 d after HFHS feeding (Supplemental Fig. 1). In contrast, HFHS feeding induced both PGC1a (↑71%) and PGC1b (↑3.2 fold). Glucocorticoid (GC) induced leucine zipper (GILZ), FK506 binding protein 5 (FKBP5), Setdb2, Mfsd2a, Lcn13, Mt1, and Mt2 are GR-target genes (113, 143) (Fig. 8). Compared to LFD-fed WT mice, WT mice fed HFHS for 15 d had marked induction of these GR-target genes. Additionally, the key gluconeogenic gene G6pc was induced 3.1 fold, and Shp tended to be higher in HFHS-fed mice (Supplemental Fig. 1). This suggests that cooperative induction of lipid catabolic genes by GR and HNF4α, rather than activation of PPARα, might mediate the early resistance to HFHS-induced fatty liver and hyperlipidemia.

### Hepatic lipid accumulation and gene dysregulation in GR knockout mice fed HFHS for 15 d

To definitively determine the role of hepatic GR in HFHS-induced lipid disorders, we specifically deleted hepatic GR in adult male mice and fed these mice with HFHS for 15 d. Histology showed that GR knockout mice had more hepatocyte vacuolization than the WT mice (Fig. 9A & 9B). Blood glucose tended to be higher in GR-knockout than WT mice (Fig. 9C). The liver/body weight ratio was 21% higher in GR knockout mice than WT mice (Fig. 9D), and hepatic levels of triglycerides and cholesterol were 95% and 56% higher than WT mice (Fig. 9E), respectively. Blood levels of triglycerides and cholesterol were similar between WT and GR knockout mice (data not shown). To understand the mechanism of aggravated lipid accumulation in HFHS-fed GR knockout mice, we used real-time PCR to determine changes in hepatic mRNA expression. As expected, hepatic expression of exon 3 of GR mRNA was remarkably decreased (↓88%), demonstrating the high efficiency of hepatic GR deletion in these inducible GR knockout mice (Fig. 9F). Consistently, GR knockout mice had marked down-regulation of GR-target genes, namely Gilz (↓69%), Mfsd2a (↓80%), Setdb2 (↓22%), Lcn13 (↓92%), and Mt1 (↓71%) (Fig. 9F). In line with a trend of higher blood glucose, hepatic G6pc tended to be higher (↑61%) in GR knockout mice (Fig. 9F). Thus, activation of GR is not responsible for hepatic induction of G6pc by HFHS (Supplemental Fig. 1). In this regard, high glucose induces glucotoxicity and G6pc expression via activating hypoxia-inducible factor (HIF)-1α or the CREB-binding protein (39). Consistent with increased hepatic lipid accumulation, GR knockout mice had induction of key lipogenic transcription factors Pparg (↑92%) and Srebp-1c (↑1.4 fold), lipogenic genes Pdk4 (↑2.7 fold), Plin2 (↑46%), and Cd36 (↑72%), as well as trend of induction of Lpl (↑50%, p = 0.07) and Fasn (↑67%, p = 0.13) (Fig. 9G). Additionally, GR knockout mice tended to have higher proinflammatory cytokines interleukin 1b (Il1b) (↑49%, p = 0.13) and C-C motif chemokine ligand 2 (Ccl2) (↑57%, p = 0.09). HLA-F-adjacent transcript 10 (FAT10) is an immune-cell-enriched ubiquitin-like modifier dramatically induced in liver injury and liver diseases (36, 37, 76). Surprisingly, GR knockout mice had 2.1 fold higher Fat10 than WT mice. Additionally, GR knockout mice had a marked 1.7 fold higher key profibrotic collagen 1a1 (Col1a1) (Fig. 9G), suggesting a role of hepatocellular GR in the protection against HFHS-induced liver fibrosis.

**Figure 9.**
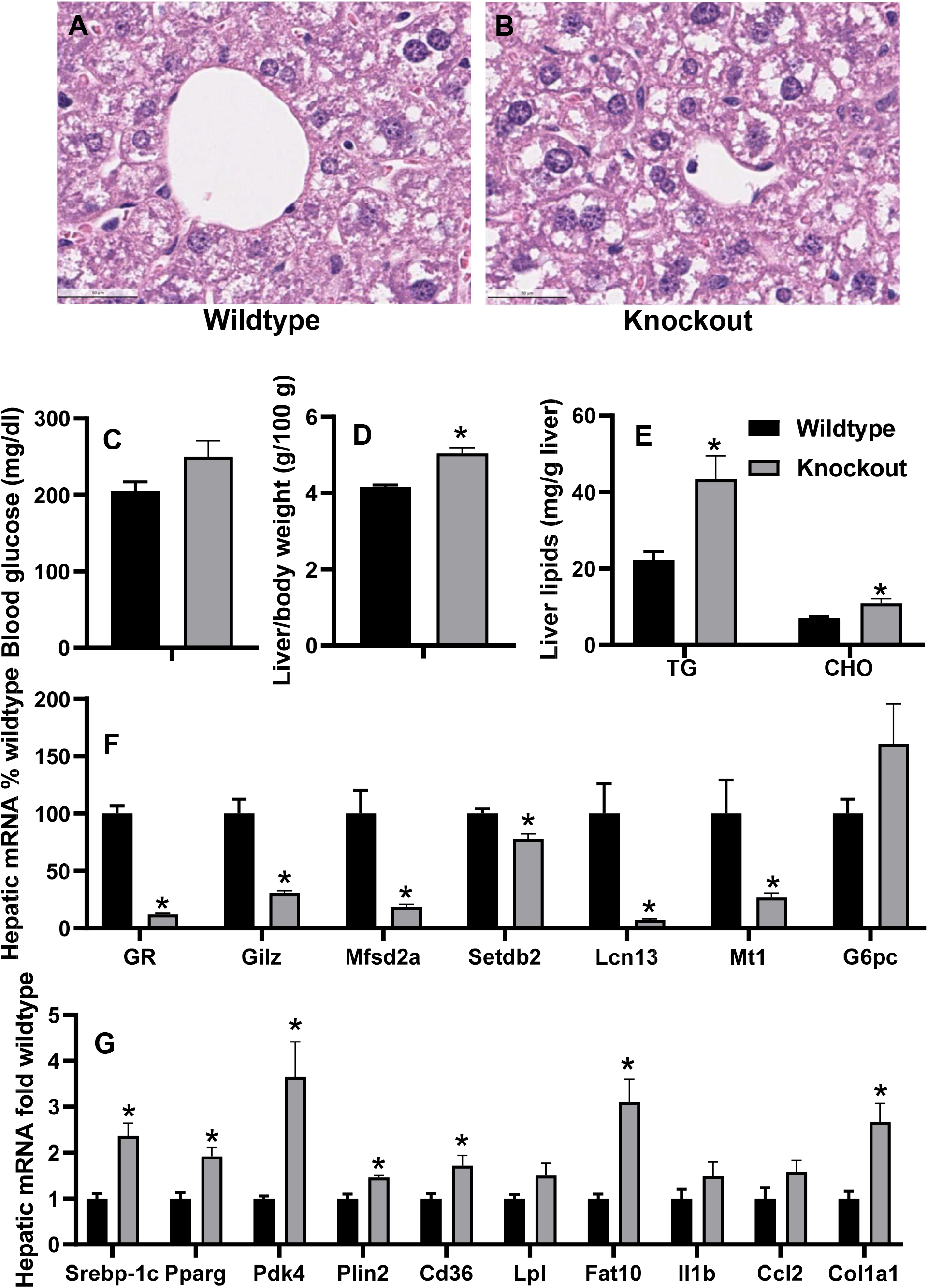
Effects of liver-specific knockout of glucocorticoid receptor (GR) on hepatic gene expression and lipid metabolism in adult male mice. (A & B) Liver histology (H&E staining, 5 μm, 400X); (C) blood glucose; (D) liver/body weight ratio; (E) hepatic lipids, and (F & G) real-time PCR quantification of hepatic mRNAs in high-fat-high-sugar (HFHS)-fed adult male GR knockout mice. N = 6 per group, mean ± SE. Hepatic mRNAs were normalized to Pgk1, with wildtype values set at 1.0 or 100%. * p < 0.05 versus wildtype control.

## Discussion

In summary, in adult male WT mice fed HFHS for 15 d, hepatic HNF4α- and GR-target lipid catabolic genes are induced, whereas lipogenic gene Scd1 were down-regulated, which is associated with their resistance to fatty liver and hyperlipidemia. In the HFHS-fed HNF4α heterozygote mice, hepatic down-regulation of HNF4α- and GR-target lipid catabolic genes and induction of lipogenic genes (e.g. Srebp-1c, Fasn, Scd1, and Acly) appear to be responsible for increases of serum and hepatic lipids. HNF4α potently inhibits the transactivation of mouse and human SREBP-1C by LXR. In contrast, down-regulation of genes important for hepatic catabolism and efflux of lipids appear to be responsible for hepatic lipid accumulation and hypolipidemia in HNF4α knockout mice. Surprisingly, hepatic nuclear GR proteins are gene-dosage-dependently decreased in HFHS-fed HNF4α heterozygote and knockout mice. In reporter assays, GR acts as a key modulator of HNF4α and PPARα to induce lipid catabolic genes and suppress lipogenic genes. Consistently, HFHS-fed GR knockout mice had increased lipid accumulation, down-regulation of lipid catabolic genes, and induction of lipogenic genes.

The present study suggests a novel key role of hepatic HNF4α in preventing not only fatty liver but also hyperlipidemia, partly via inhibiting the induction of master lipogenic factor SREBP-1C by LXR. When activated by oxysterols or insulin, LXR bind to LXR response element (LXRE) to transactivate genes. Activation of LXR in liver causes fatty liver and hyperlipidemia due to increased *de novo* lipogenesis via induction of SREBP-1C (63). Unsaturated FAs potently down-regulate SREBP-1C to feedback inhibit lipogenesis (45); how such feedback regulation is compromised in steatosis is poorly understood. Our data indicate that HNF4α antagonizes the transactivation of SREBP-1C by LXR to control hepatic lipid metabolism, and thus establish deficiency of HNF4α as a key mechanism of the dysregulation of the two master lipogenic factors LXR and SREBP-1C during hepatosteatosis and hyperlipidemia.

HNF4α deficiency causes dramatic up- and down-regulation of a very large number of genes. Despite extensive studies of how HNF4α transactivates its target genes, little is known how HNF4α suppresses gene expression. The corepressor SHP can inhibit the transcriptional activities of a large number of transcription factors. Interestingly, HNF4α has been shown to play a critical role in determining the cellular localization of SHP: SHP remains in the cytosol in the absence of HNF4α (154). We found that SHP mediates the inhibition of LXR-activation of SREBP-1C promoter by HNF4α. SHP promotes early fatty liver by inducing PPARγ, whereas loss of SHP aggravates hepatic inflammation and liver cancer (83, 152). Interestingly, human SHP promoter is stimulated by HNF4α (72). The role of SHP in mediating the inhibitory effects of HNF4α on hepatic lipogenesis, inflammation, and liver cancer warrants further investigation.

The present study demonstrates that partial loss of HNF4α results in a moderate elevation of HDL cholesterol and a more marked elevation of LDL/VLDL cholesterol, resulting in an elevated total/HDL cholesterol ratio (Fig. 1) and CAD risk. Liver-specific overexpression of endothelial lipase (LIPG) dramatically decreases blood levels of total and HDL-cholesterol by accelerating HDL catabolism and increasing hepatic uptake of HDL-cholesteryl ether (129). We found a novel gene-dosage-dependent critical role of HNF4α in determining hepatic expression of LIPG in HFHS-fed mice (Fig. 5D). Hepatic expression of LIPC and SR-BI (other genes in HDL-cholesterol uptake) as well as ABCA1, APOA1, and APOA2 (key genes in HDL biogenesis) remain unchanged in the heterozygote mice (Fig. 5D, 5E), strongly suggesting that hepatic down-regulation of LIPG and the resultant decreased hepatic uptake of HDL-cholesterol may be responsible for the elevated HDL-cholesterol in the HNF4α heterozygote mice. Very interestingly, in an animal model of atherosclerosis, LIPG expression is enhanced in the aorta and reduced in the liver of mice developing atherosclerosis (149). In addition to regulating LIPG-mediated HDL-cholesterol uptake, the present study shows that HNF4α regulates hepatic efflux of lipid and cholesterol by differentially affecting APOC3 and APOA4 expression. In addition to APOB, APOC3 and APOA4 are two of the major apolipoproteins responsible for hepatic secretion of VLDL (137). In contrast to the association of APOC3 with the small and dense VLDL (124), APOA4 promotes the expansion of larger VLDL proteins that are thought to have less CVD risk (137). Interestingly, lower serum APOA4 but higher serum APOC3 are associated with increased risk of CAD (67, 99). Similar to CAD patients, Apoa4 was lower, whereas Apoc3 tended to be higher, in HFHS-fed HNF4α heterozygote mice than WT mice (Fig. 5D). Taken together, our data strongly suggest that partial HNF4α deficiency may be a key driver of not only NAFLD, but also hyperlipidemia and CAD risk during HFHS intake.

The present study discovers novel critical roles of HNF4α heteroinsufficiency and knockout in regulating hepatic CYP7A1 expression and BA metabolism during HFHS intake. In the knockout mice, the dramatic down-regulation of CYP7A1, CYP8B1, and CYP2C70 but a moderate down-regulation of CYP27A1 (59%) will shift the BA biosynthesis from CA to CDCA, resulting in a trend of decreased total CA (↓34%) but a marked 4.9 fold increase of total CDCA. In contrast, the dramatic down-regulation of BAAT is likely responsible for the marked increase of free DCA and decrease of T-DCA in the knockout mice. In HNF4α knockout mice, the much less decreases in total T-BAs than total sulfated BAs, despite similarly dramatic down-regulation of the conjugating enzymes BAAT and SULT2A8, is likely due to the fact that most of the sulfated BAs are excreted in the urine without recycling (133), whereas the majority of T-BAs are recycled via enterohepatic circulation. In contrast to the moderate decreases of T-BAs and dramatic decreases of G-BAs in the liver, a previous study indicates that the gallbladder bile of HNF4α knockout mice have dramatic increases in free BAs (>100 fold) as well as G-DCA and G-CDCA (>20 fold) but only moderate increases of the major taurine-conjugated BAs T-CA, T-MCA, and T-DCA (54). Thus, HNF4α deficiency leads to increased basolateral efflux of sulfated BAs for urinary excretion and increased canalicular efflux of free as well as taurine- and glycine-conjugated BAs into the bile to alleviate hepatocyte accumulation of BAs and the resultant liver injury. Currently, the role of CYP7A1 in HFD-induced NAFLD is controversial (18, 33). Mouse Cyp7a1 has LXR site and is induced by cholesterol. In contrast, the lack of LXR site in human CYP7A1 promoter prevents hepatic induction of human CYP7A1 by high cholesterol, resulting in increased hypercholesterolemia when fed a HFD (18). Transgenic mice overexpressing CYP7A1 have expanded BA pool and are resistant to HFD-induced obesity, fatty liver, and insulin resistance (74). In contrast, Cyp7a1 knockout mice are also resistant to HFD-induced obesity, fatty liver, and insulin resistance due to induction of the alternative pathway Cyp27a1 and Cyp7b1 and more production of the hydrophilic MCA (33). However, humans lack the CYP2C70-catalyzed CDCA-MCA pathway and thus have a much more hydrophobic BA profile than rodents (128). Consequently, CYP7A1 mutation in humans causes hypercholesterolemia, hypertriglyceridemia, and accumulation of cholesterol in the liver, despite induction of the alternative pathway (106). Thus, the more hydrophilic BAs in livers of HNF4α heterozygote mice, despite a marked down-regulation of Cyp7a1, is likely a rodent-specific phenotype. Humans with hepatic HNF4α deficiency may experience more marked disorder in cholesterol, BA, and lipid metabolism when on HFD than the HFD-fed HNF4α heterozygote mice.

The present study suggests that partial and total deficiency of HNF4α may differentially modulate the BA receptor FXR signaling in the liver. Within the cells, free and taurine-conjugated CA and CDCA have similar potency in activating FXR (139), whereas T-MCAs are FXR antagonists (116). The marked increase of total CDCA, the most potent FXR agonist in HNF4α knockout mice (Fig. 4D) is consistent with the previous report of activation of FXR in these mice (134). In contrast, the increase of the FXR antagonist T-MCA and trend of decreases in the FXR agonist DCA and CA in livers of HNF4α heterozygote mice (Fig. 4E) might result in impaired FXR signaling. It is noteworthy that although the FXR-target gene SHP tended to be induced in both the HNF4α heterozygote and knockout mice (Fig. 5B), SHP can also be induced by other nuclear receptor(s). Particularly, LXRα can induce human SHP by binding to the DNA response element in the SHP promoter that overlaps with the previously characterized bile acid response element (41), and our data mining found that LXRα also tends to induce Shp in mouse liver (GSE2644).

Literature suggests that activation of GR in extrahepatic tissues worsens NAFLD, whereas hepatic GR may be protective in lipid metabolism during HFD intake (6, 86, 91, 103, 123). Although many acute effects of GCs mobilize energy and cause weight loss, chronically elevated circulating GCs motivate people to overeat HFHS food and promote obesity and NAFLD in extrahepatic tissues (6, 123). Importantly, circulating GCs markedly increase in mice with liver-specific GR deficiency (91), and literature support protective role of hepatic GR against fatty liver during high FA flux (86, 91, 103). In this regard, Stat5/GR double knockout mice have more severe steatohepatitis than Stat5 knockout mice (91). Liver-specific overexpression of HSD11B1 in HFD-fed mice down-regulates LXRα and tends to ameliorate steatosis (103). Importantly, on a HFD, GR +/− mice have higher hepatic TG than WT mice despite similar feed intake (86). In NAFLD patients, hepatic activities of HSD11B1 decrease, whereas the cortisol-inactivation enzyme increase (3, 64), which may result in local GC deficiency and concomitant activation of the hypothalamic–pituitary–adrenal (HPA) axis to trigger GC release (3, 64). In contrast, a specific HSD11B1 inhibitor worsens liver fibrosis in mice (158). Our results from HFHS-fed mice with adult liver-specific knockout of GR provide definitive evidence that hepatocellular GR is essential in the protection against HFHS-induced fatty liver, and hepatic GR may also be important in preventing the development of liver fibrosis.

The present study discovers novel GR-target genes important in the regulation of lipid metabolism. In addition to the known GR-target genes GILZ, MT1, and SETDB2, we identified five novel GR-target genes, namely SREBP-1C, PPARγ, LCN13, MFSD2A, and FAT10 (Fig. 9). Induction of the two key lipogenic transcription factors SREBP-1C and PPARγ may be responsible for hepatic induction of lipgenic genes and increased lipid accumulation in the GR knockout mice. Currently, the mechanism of hepatic suppression of the lipogenic PPARγ by GR is unknown. SREBP-1C is a transactivator (29), whereas GILZ is a transrepressor of PPARγ (122). Thus, derepression of SREBP-1C or down-regulation of GILZ may lead to the induction of PPARγ in GR knockout mice. In hepatocytes, LCN13 directly suppresses hepatic gluconeogenesis and lipogenesis but increases fatty acid β oxidation (155). Currently, the role of MFSD2A, the key transporter for ω3-fatty acids in fatty liver is still unclear (8). In view of the important role of essential ω3-fatty acids in the protection against fatty liver, the potential role of GR-MFSD2A pathway in regulating hepatic lipid metabolism warrants further investigation. FAT10 is a 165-amino acid ubiquitin-like protein for ubiquitin-independent proteasomal degradation (4). GR antagonizes STAT3 and NF-kB (68). FAT10 is synergistically induced by NF-kB and STAT3, and FAT10 mediates NF-KB activation (5, 20, 117). Interestingly, FAT10-null mice have increased insulin sensitivity and fatty acid oxidation and decreased fat mass (12). The role of GR/FAT10 pathway in regulating hepatic protein homeostasis and steatohepatitis warrants further investigation.

The present study demonstrates that hepatic GR has an important role in modulating the transcriptional activity of HNF4α and PPARα during hepatosteatosis. GR and HNF4α cooperatively induce genes that promote lipid catabolism, such as Cyp7a1, Por, SetDb2, and Lcn13, whereas GR antagonizes the activation of the lipogenic Apoc3 by HNF4α. In parallel, treatment of primary human hepatocytes with the GR agonist dexamethasone increases the expression of CYP7A1 and production of CA (90). Activation of the FA receptor PPARα promotes FA catabolism to protect against fatty liver during fasting and HFD intake (88). However, PPARα can also induce key lipogenic genes (27, 32) and enhance lipogenesis, which may abate its beneficial effects on lipid catabolism. Particularly, PPARα induces the key lipogenic transcription factor Srebp-1c by directly binding to its promoter (32), and induction of lipogenic genes by chronic PPARα activation is abolished in Srebp1 knockout mice (101). Consequently, both protective and aggravating effects of PPARα activation on fatty liver in mice have been reported (27, 88, 147). Interestingly, GR agonist can antagonize the induction of lipogenic genes by PPARα agonist in mouse hepatocytes (108). The present study demonstrates that GR antagonizes the induction of Srebp-1c, PLIN2, and CD36 by PPARα in reporter assay, and the induction of Srebp-1c, Plin2, Cd36, and Pdk4 are associated with increased lipid accumulation in HFHS-fed GR knockout mice. Thus, hepatic GR signaling might be a key determinant of the beneficial versus detrimental effects of PPARα activation on hepatic lipid metabolism, which warrants further investigation. These lipogenic genes Plin2, Pdk4, and Cd36 were also highly elevated in HFHS-fed HNF4α knockout mice (Fig. 5). AMPK is a key modulator of GR transcriptional activity (93, 108). When AMPK activity is low, GR agonist potently inhibits the induction of Pdk4 by PPARα (108). Importantly, AMPK activity is markedly decreased in HNF4α knockout mice (Fig. 6). Thus, impairment of GR signaling in HNF4α knockout mice, likely due to the loss of AMPK-mediated GR phosphorylation (93), may contribute to the super-induction of PPARα-target lipogenic genes Plin2, Pdk4, and Cd36. Further in-depth study of liver-specific GR-HNF4α and GR-PPARα/PPARγ crosstalk will help understand not only fatty liver, but also the progression from simple steatosis to steatohepatitis and cirrhosis.

In conclusion, the present study clearly demonstrates that HNF4α and GR play key roles in the protection against HFHS-induced elevation of circulating and hepatic lipids. When the activation of PPARα signaling is impaired by fructose overdose, the cooperative induction of lipid catabolic genes and suppression of lipogenic genes by HNF4α and GR, modulated by AMPK, may play a key role in the early resistance to HFHS-induced fatty liver and hyperlipidemia (Graphic abstract). The gene-environment interaction between an obesogenic environment and genetic susceptibility drives the pathogenesis of metabolic disease and NAFLD. Numerous pathological mutations with decreased GC sensitivity have been identified in the human GR gene (136). Hepatic HNF4α expression is highly variable in humans (144), and is markedly decreases in diabetes and NAFLD (73, 145, 146). Very interestingly, obesity and elevated blood triglycerides is associated with a high frequency of missense T130I mutation of HNF4α in non-diabetic indigenous Mexicans with high intake of HFHS (42). Partial deficiency of HNF4α and/or GR due to genetic polymorphism and/or metabolic stresses is thus likely a key mechanism of the loss of resistance to hepatosteatosis and hyperlipidemia during HFHS intake. The identification of partial deficiency of HNF4α and GR as potential key drivers of NAFLD and hyperlipidemia may help develop novel pharmaceutical and dietary interventions for HFHS-induced NAFLD and CAD. Although the activating ligand for HNF4α has not been identified, small activating RNA for HNF4α has been used to successfully induce HNF4α and ameliorate hyperlipidemia and fatty liver in HFD-fed rats (52). In view of GC’s potent anti-inflammatory effects, liver-specific activation of GR may protect against HFHS-induced NAFLD and the progression from NAFLD to NASH and cirrhosis.

## Experimental procedures

### Generation and treatment of HNF4α heterozygote and knockout mice

We generated tamoxifen-inducible knockout mice by crossing HNF4α floxed mice (82) with mice carrying tamoxifen-inducible estrogen-receptor-fused Cre under the control of an albumin promoter (SA^+/CreERT2^) (10). Adult (8-week old) male WT (Cre/−), heterozygote (HNF4α fl/+, Cre/+), and knockout (HNF4α fl/fl, Cre/+) mice (N=3-6 per group) were administered tamoxifen (T5648, Sigma, 5 mg/kg ip in corn oil) once daily for 2 d. The day after the 2nd tamoxifen treatment, mice were fed a (A) LFD (10% fat kcal, D12450B, Research Diets, Inc) or (B) HFHS that contained high-fat diet (HFD, 60% fat kcal, #D12492, Research Diets) and a high-sugar drinking water with 42 g/L of sugar (55% fructose/45% sucrose) (62). All mice were allowed water and feed *ad libitum* and sacrificed 15 d or 6 weeks after diet treatments without pre-fasting to collect liver and blood samples. Liver tissues were snap frozen in liquid nitrogen upon collection and stored at −80 °C until use. To prepare serum samples, the clotted blood samples were centrifuged at 8000 rpm for 10 min. All animals received humane care and all animal procedures in this study were approved by the Institutional Animal Care and Use Committee (IACUC) of the SUNY Upstate Medical University.

### Generation and treatment of GR knockout mice

We generated mice with tamoxifen-inducible liver-specific knockout of GR by crossing GR floxed mice (Stock# 021021, Jackson Laboratory) (87) with the SA^+/CreERT2^ mice (10). Adult (8-10 week old) male WT (GR fl/fl, Cre/−) and GR knockout (GR fl/fl, Cre/+) mice (N=6 per group) were administered tamoxifen (T5648, Sigma, 5 mg/kg ip in corn oil) once daily for 2 d. The day after the 2nd tamoxifen treatment, mice were fed HFHS (62) and sacrificed 15 d after HFHS feeding, without pre-fasting to collect liver and blood samples. Blood glucose was determined by Care Touch Blood Glucose Meter. Liver tissues were snap frozen in liquid nitrogen upon collection and stored at −80 °C until use. To prepare serum samples, the clotted blood samples were centrifuged at 8000 rpm for 10 min.

### Construction of reporter and expression vectors

The PCR primers were synthesized by Integrated DNA Technologies (IDT). Total genomic DNA extracted from liver mouse and HEK293 cells were used as templates for PCR-cloning of the mouse/human promoter/intron fragments into the KpnI/MluI sites of pGL3 basic luciferase reporter vector (Promega). The glutamic acid (E) to lysine (K) at 363 (E363K) mutant of HNF4A2 was generated using Q5® Site-Directed Mutagenesis Kit (New England Biolabs). The pcDNA3-SHP-Flag vector for SHP was synthesized by GenScript. The primers and sequence information of all reporter constructs is provided in Supplemental table S1 (https://figshare.com/s/650749d0f09c1c5d4b2e). All the constructed vectors were verified by sequencing.

### Transient transfection and dual-luciferase assay

HEK293 cells were cultured with MEM medium (Corning) supplemented with 10% fetal calf serum. Twenty-four hours after seeding, transfection was conducted using Lipofectamine 2000 (Invitrogen), following the manufacturer's protocol. In the 96-well-plate, each well *was* transfected with firefly luciferase vectors, the control renilla luciferase vector pRL-CMV, and/or the mammalian expression vectors for human HNF4A2 (#31100, Addgene), LXRα (110), GR (#15534, Addgene), SHP, and/or the pCMX backbone vector to add up the total DNA vectors to 100 ng. Dexamesasone (10 nM) was added 1 h after cells were transfected with the GR expression vector. Twenty-four hours after transfection, cells were harvested for dual-luciferase assay using Dual-Glo™ luciferase assay system (Promega) and GloMax Luminometer (Promega), following the manufacturer’s protocol. The ratios of firefly/renilla luciferase activities were calculated as the normalized reporter activity, with the control values set at 1.0.

### Western blot

Proteins in cytosolic or nuclear extracts from mouse livers were resolved in sodium dodecyl sulphate-polyacrylamide gel electrophoresis. Western blot quantification of proteins was conducted with primary antibodies as follows: HNF4α (#3113, Cell Signaling), GR (sc-1002, Santa Cruz), β-catenin (#9562, Cell Signaling), acetyl-CoA carboxylase (ACC, #3676, Cell Signaling), phospho-ACC (#11818, Cell Signaling), phospho-AMP-activated protein kinase α (AMPKα) (Thr172) (#2535, Cell Signaling), fatty acid synthase (Fasn, # 3180, Cell Signaling), Srebp-1c (SAB4502850, Sigma), histone H3 (#4499, Cell Signaling), and β-actin (#5779-1, Epitomics). Primary antibodies were revealed with HRP-conjugated secondary antibodies (Anti-mouse IgG, #7076; Anti-rabbit IgG, #7074, Cell Signaling) and ECL Western Blotting Substrate (W1015, Promega). ChemiDoc™ XRS+System (Bio-Rad) and Image-J software were used for band capturing and density analysis. Nuclear and cytosolic levels of proteins were normalized to histone H3 and β-actin, respectively, with the wildtype values set as 1.0.

### Real-time PCR

Total RNAs from mouse livers were isolated by RNA-STAT60 (Tel-Test) and quantified by NanoDrop^™^ spectrophotometer (Thermo Scientific). For real-time PCR, 1 μg of RNA was reverse transcribed using the High-Capacity RNA-to-cDNA^™^ Kit (Applied Biosystems®, life technologies) for cDNA synthesis, following the manufacturer's instructions. iQ^™^ SYBR® Green Supermix (Bio-Rad) was applied to quantify mRNAs using MyiQ2^™^ Two-Color Real-Time PCR Detection System (Bio-Rad). Alternatively, total RNAs from mouse livers were isolated by RNA-STAT60 (Tel-Test) and quantified by NanoDrop^™^. RNA was reverse transcribed using iScript^™^ cDNA Synthesis Kit (Bio-Rad), following which qPCR was performed using iTaq^™^ Universal SYBR® Green Supermix (Bio-Rad) using Bio-Rad’s CFX Maestro (Version: 4.1.2433.1219.). The amounts of mRNA were calculated using the comparative CT method, which determines the amount of target gene normalized to peroxiredoxin 1 (Prdx1) or phosphoglycerate kinase 1 (Pgk1), two of the most stable housekeeping genes (30, 153). The specificity of the real-time PCR primers was verified using the no-reverse-transcriptase control. Sequences of real-time PCR primers (synthesized by IDT) were listed in Supplemental table S2 (https://figshare.com/s/52b2fd910184cc82d322).

### Determination of lipids in mouse liver

Frozen liver tissue were homogenized in buffer containing 18 mM Tris, pH 7.5, 300 mM mannitol, 10 mM EGTA, and 0.1 mM phenylmethylsulfonyl fluoride (81). Liver homogenates were mixed with chloroform:methanol (2:1) and incubated overnight at room temperature with occasional shaking. The next day, H_2_O was added, vortexed, and centrifuged at 3000 *g* for 5 min. The lower lipid phase was collected and concentrated by speed vacuum concentrator. The lipid pellets were dissolved in a mixture of 270 μl of isopropanol and 30 μl of Triton X-100 to determine triglycerides and total cholesterol using commercial analytical kits (Pointe Scientific, Inc, Canton MI).

### Determination of blood levels of glucose, insulin, cholesterol, free fatty acids, and ketone bodies

Blood levels of glucose were quantified with Precision Xtra Glucose Monitor (#179837, Bound Tree Medical, LLC, Chicago, IL). Serum levels of insulin were determined with an ELISA kit (#90080, Crystal Chem USA, Downers Grove, IL). Serum levels of total cholesterol, HDL cholesterol, and LDL/VLDL cholesterol were quantified with HDL and LDL/VLDL assay kit (EHDL-100, Bioassay Systems). Serum levels of free fatty acids were analyzed with EnzyChrom™ Free Fatty Acid Assay Kit (EFFA-100, Bioassay Systems). Serum levels of β-hydroxybutyrate were quantified with a colorimetric assay kit (#700190, Cayman Chemical, Ann Arbor, MI).

### Bile acid quantification

Bile acids in liver and serum in LFD-fed wild-type mice and HFHS-fed wild-type, HNF4α heterozygote, and HNF4α knockout mice were quantified by liquid chromatography-tandem mass spectrometry (LC–MS/MS), as we described previously with some modifications (Bathena et al., 2013; Huang, Bathena, Csanaky, & Alnouti, 2011). Briefly, a Waters ACQUITY ultra‐performance LC system (Waters, Milford, MA, USA) coupled to a 4000 Q TRAP® quadrupole linear ion trap hybrid MS with an electrospray ionization source (Applied Biosystems, MDS Sciex, Foster City, CA, USA) was used. The following MS source settings were used: ion spray voltage, −4500 V; temperature, 550°C; curtain gas, 10; gas‐1, 40; gas‐2 40 (arbitrary units); collision gas pressure, high; Q1/Q3 resolution, unit; and interface heater, on. Mobile phase consisted of 7.5 mM ammonium bicarbonate, adjusted to pH 9.0 using ammonium hydroxide (mobile phase A) and 30% acetonitrile in methanol (mobile phase B) at a total flow rate of 0.2 ml min^−1^. The gradient profile was held at 52.5% mobile phase B for 12.75 minutes, increased linearly to 68% in 0.25 minutes, held at 68% for 8.75 minutes, increased linearly to 90% in 0.25 minutes, held at 90% for 1 minute and finally brought back to 52.5% in 0.25 minutes followed by 4.75 minutes re‐equilibration (total run time of 28 minutes per sample). For the preparation of calibration curves, blank matrices were obtained by charcoal stripping as described previously (Bathena et al., 2013; Huang, Bathena, Csanaky, & Alnouti, 2011). Eleven‐point calibration curves were prepared by spiking 10 μl of appropriate standard solution into 50 μl stripped serum and 100 μl stripped liver homogenate at final concentrations ranging from 1 to 1000 ng ml^−1^.

For the preparation of serum samples, 50 μl of serum samples were spiked with 10 μl of internal standard, 1 ml of ice‐cold alkaline acetonitrile (5% NH_4_OH) was added and samples were vortexed. Samples were then centrifuged at 16 000 *g* for 10 minutes and the supernatants were aspirated, evaporated under vacuum and reconstituted in 100 μl of 50% MeOH solution. For liver samples, approximately 100 mg of liver was homogenized in 4 volumes of water. A 100 μl of liver homogenate was spiked with 10 μl IS and 2ml of ice-cold alkaline ACN was added. Samples were vortexed, and shaken continuously for 15 min, and then centrifuged at 16,000×g for 10 min. The supernatant was aspirated and the pellet was extracted with another 1ml of ice-cold alkaline ACN. Supernatants from the 2 extraction steps were pooled, evaporated, and reconstituted in a 100μl of 50% MeOH.

The hydrophobicity index (HI) of bile acids was calculated by the method of Heuman (48).

### Statistical analysis

All values were expressed as mean ± S.E. For comparison of two groups, the two-tailed student’s t-test was used to determine the statistical difference, which was set at p < 0.05. For multiple comparisons, analysis of variance (ANOVA) was performed, followed by the Student-Newman-Keuls Method in SigmaPlot 12.5, with significance set at p < 0.05.

## Footnotes

Funding was provided partly by grants to H.L. from the National Institute of Health (R01ES019487, R21AA027349, and R03CA241781).

## Acknowledgements

We would like to thank Dr. Pierre Chambon (IGBMC, Illkirch, France) for providing the SA^+/CreERT2^ mice (118), Dr. David Mangelsdorf for providing the LXR expression vector and the reporter vector for mouse Srebp-1c promoter, and Dr. Etienne Lefai for providing the reporter vector for human SREBP-1C promoter (25, 110).

## Authors contributions

HL designed the experiments, performed real-time PCR analysis, and wrote the manuscript. XL performed animal and cellular experiments and analyzed the results. SG performed Western blot experiments and analyzed the results. XL and RW performed hepatic lipid assay and real-time PCR experiments and analyzed the results. SJ revised critically for important intellectual content. DK and WL conducted LC-MS/MS analysis of serum and hepatic bile acids and analyzed the results. YA analyzed the results of bile acids analysis and edited the manuscript.

---

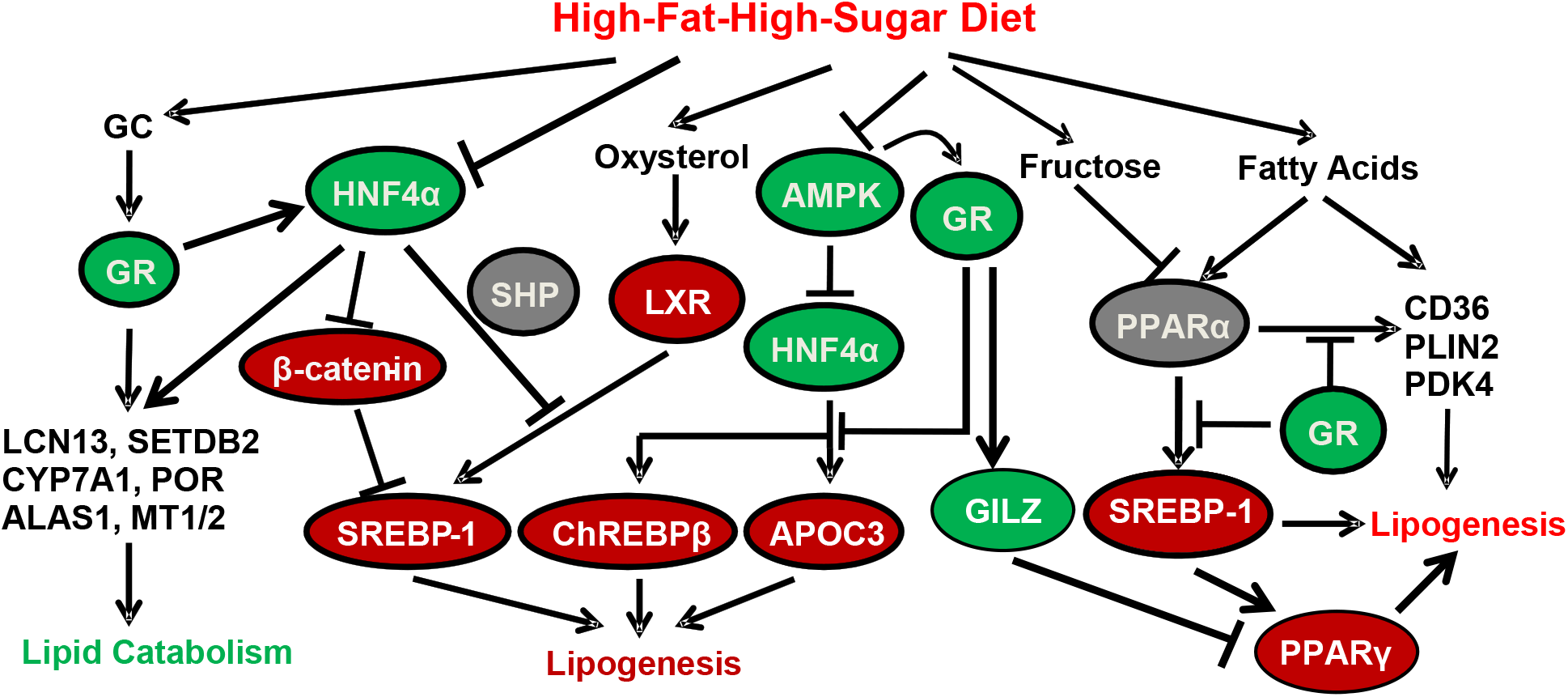
Graphic abstract.

## References

1. Abiola M, Favier M, Christodoulou-Vafeiadou E, Pichard AL, Martelly I, and Guillet-Deniau I. Activation of Wnt/beta-catenin signaling increases insulin sensitivity through a reciprocal regulation of Wnt10b and SREBP-1c in skeletal muscle cells. PloS one 4: e8509, 2009.

2. Adams JM, and Jafar-Nejad H. The Roles of Notch Signaling in Liver Development and Disease. Biomolecules 9: 2019.

3. Ahmed A, Rabbitt E, Brady T, Brown C, Guest P, Bujalska IJ, Doig C, Newsome PN, Hubscher S, Elias E, Adams DH, Tomlinson JW, and Stewart PM. A switch in hepatic cortisol metabolism across the spectrum of non alcoholic fatty liver disease. PloS one 7: e29531, 2012.

4. Aichem A, and Groettrup M. The ubiquitin-like modifier FAT10 in cancer development. The international journal of biochemistry & cell biology 79: 451–461, 2016.

5. Basler M, Buerger S, and Groettrup M. The ubiquitin-like modifier FAT10 in antigen processing and antimicrobial defense. Molecular immunology 68: 129–132, 2015.

6. Bazhan N, and Zelena D. Food-intake regulation during stress by the hypothalamo-pituitary-adrenal axis. Brain research bulletin 95: 46–53, 2013.

7. Bellentani S, Scaglioni F, Marino M, and Bedogni G. Epidemiology of non-alcoholic fatty liver disease. Dig Dis 28: 155–161, 2010.

8. Berger JH, Charron MJ, and Silver DL. Major facilitator superfamily domain-containing protein 2a (MFSD2A) has roles in body growth, motor function, and lipid metabolism. PloS one 7: e50629, 2012.

9. Besse A, Wu P, Bruni F, Donti T, Graham BH, Craigen WJ, McFarland R, Moretti P, Lalani S, Scott KL, Taylor RW, and Bonnen PE. The GABA transaminase, ABAT, is essential for mitochondrial nucleoside metabolism. Cell metabolism 21: 417–427, 2015.

10. Bonzo JA, Ferry CH, Matsubara T, Kim JH, and Gonzalez FJ. Suppression of hepatocyte proliferation by hepatocyte nuclear factor 4alpha in adult mice. The Journal of biological chemistry 287: 7345–7356, 2012.

11. Brown MS, and Goldstein JL. A proteolytic pathway that controls the cholesterol content of membranes, cells, and blood. Proceedings of the National Academy of Sciences of the United States of America 96: 11041–11048, 1999.

12. Canaan A, DeFuria J, Perelman E, Schultz V, Seay M, Tuck D, Flavell RA, Snyder MP, Obin MS, and Weissman SM. Extended lifespan and reduced adiposity in mice lacking the FAT10 gene. Proceedings of the National Academy of Sciences of the United States of America 111: 5313–5318, 2014.

13. Carr RM, and Ahima RS. Pathophysiology of lipid droplet proteins in liver diseases. Experimental cell research 340: 187–192, 2016.

14. Chang B, Xu MJ, Zhou Z, Cai Y, Li M, Wang W, Feng D, Bertola A, Wang H, Kunos G, and Gao B. Short- or long-term high-fat diet feeding plus acute ethanol binge synergistically induce acute liver injury in mice: an important role for CXCL1. Hepatology 62: 1070–1085, 2015.

15. Chapman K, Holmes M, and Seckl J. 11beta-hydroxysteroid dehydrogenases: intracellular gate-keepers of tissue glucocorticoid action. Physiological reviews 93: 1139–1206, 2013.

16. Chella Krishnan K, Floyd RR, Sabir S, Jayasekera DW, Leon-Mimila PV, Jones AE, Cortez AA, Shravah V, Peterfy M, Stiles L, Canizales-Quinteros S, Divakaruni AS, Huertas-Vazquez A, and Lusis AJ. Liver Pyruvate Kinase Promotes NAFLD/NASH in Both Mice and Humans in a Sex-Specific Manner. Cellular and molecular gastroenterology and hepatology 2020.

17. Chen J, Zhao KN, and Chen C. The role of CYP3A4 in the biotransformation of bile acids and therapeutic implication for cholestasis. Ann Transl Med 2: 7, 2014.

18. Chen JY, Levy-Wilson B, Goodart S, and Cooper AD. Mice expressing the human CYP7A1 gene in the mouse CYP7A1 knock-out background lack induction of CYP7A1 expression by cholesterol feeding and have increased hypercholesterolemia when fed a high fat diet. The Journal of biological chemistry 277: 42588–42595, 2002.

19. Chiu HK, Qian K, Ogimoto K, Morton GJ, Wisse BE, Agrawal N, McDonald TO, Schwartz MW, and Dichek HL. Mice lacking hepatic lipase are lean and protected against diet-induced obesity and hepatic steatosis. Endocrinology 151: 993–1001, 2010.

20. Choi Y, Kim JK, and Yoo JY. NFkappaB and STAT3 synergistically activate the expression of FAT10, a gene counteracting the tumor suppressor p53. Molecular oncology 8: 642–655, 2014.

21. Chypre M, Zaidi N, and Smans K. ATP-citrate lyase: a mini-review. Biochemical and biophysical research communications 422: 1–4, 2012.

22. Coffinier C, Gresh L, Fiette L, Tronche F, Schutz G, Babinet C, Pontoglio M, Yaniv M, and Barra J. Bile system morphogenesis defects and liver dysfunction upon targeted deletion of HNF1beta. Development 129: 1829–1838, 2002.

23. Csanaky IL, Lu H, Zhang Y, Ogura K, Choudhuri S, and Klaassen CD. Organic anion-transporting polypeptide 1b2 (Oatp1b2) is important for the hepatic uptake of unconjugated bile acids: Studies in Oatp1b2-null mice. Hepatology 53: 272–281, 2011.

24. Dallman MF, Pecoraro NC, and la Fleur SE. Chronic stress and comfort foods: self-medication and abdominal obesity. Brain Behav Immun 19: 275–280, 2005.

25. Dif N, Euthine V, Gonnet E, Laville M, Vidal H, and Lefai E. Insulin activates human sterol-regulatory-element-binding protein-1c (SREBP-1c) promoter through SRE motifs. The Biochemical journal 400: 179–188, 2006.

26. Dwyer BJ, Olynyk JK, Ramm GA, and Tirnitz-Parker JE. TWEAK and LTbeta Signaling during Chronic Liver Disease. Frontiers in immunology 5: 39, 2014.

27. Edvardsson U, Ljungberg A, Linden D, William-Olsson L, Peilot-Sjogren H, Ahnmark A, and Oscarsson J. PPARalpha activation increases triglyceride mass and adipose differentiation-related protein in hepatocytes. Journal of lipid research 47: 329–340, 2006.

28. Eslam M, Sanyal AJ, George J, and an international consensus p. MAFLD: A consensus-driven proposed nomenclature for metabolic associated fatty liver disease. Gastroenterology 2020.

29. Fajas L, Schoonjans K, Gelman L, Kim JB, Najib J, Martin G, Fruchart JC, Briggs M, Spiegelman BM, and Auwerx J. Regulation of peroxisome proliferator-activated receptor gamma expression by adipocyte differentiation and determination factor 1/sterol regulatory element binding protein 1: implications for adipocyte differentiation and metabolism. Molecular and cellular biology 19: 5495–5503, 1999.

30. Falkenberg VR, Whistler T, Murray JR, Unger ER, and Rajeevan MS. Identification of Phosphoglycerate Kinase 1 (PGK1) as a reference gene for quantitative gene expression measurements in human blood RNA. BMC research notes 4: 324, 2011.

31. Feng L, Yuen YL, Xu J, Liu X, Chan MY, Wang K, Fong WP, Cheung WT, and Lee SS. Identification and characterization of a novel PPARalpha-regulated and 7alpha-hydroxyl bile acid-preferring cytosolic sulfotransferase mL-STL (Sult2a8). Journal of lipid research 58: 1114–1131, 2017.

32. Fernandez-Alvarez A, Alvarez MS, Gonzalez R, Cucarella C, Muntane J, and Casado M. Human SREBP1c expression in liver is directly regulated by peroxisome proliferator-activated receptor alpha (PPARalpha). The Journal of biological chemistry 286: 21466–21477, 2011.

33. Ferrell JM, Boehme S, Li F, and Chiang JY. Cholesterol 7alpha-hydroxylase-deficient mice are protected from high-fat/high-cholesterol diet-induced metabolic disorders. Journal of lipid research 57: 1144–1154, 2016.

34. Florentino RM, Fraunhoffer NA, Morita K, Takeishi K, Ostrowska A, Achreja A, Animasahun O, Haep N, Arazov S, Agarwal N, Collin de l’Hortet A, Guzman-Lepe J, Tafaleng EN, Mukherjee A, Troy K, Banerjee S, Paranjpe S, Michalopoulos GK, Bell A, Nagrath D, Hainer SJ, Fox IJ, and Soto-Gutierrez A. Cellular Location of HNF4alpha is Linked With Terminal Liver Failure in Humans. Hepatology communications 4: 859–875, 2020.

35. Franklin MP, Sathyanarayan A, and Mashek DG. Acyl-CoA Thioesterase 1 (ACOT1) Regulates PPARalpha to Couple Fatty Acid Flux With Oxidative Capacity During Fasting. Diabetes 66: 2112–2123, 2017.

36. French BA, Oliva J, Bardag-Gorce F, and French SW. The immunoproteasome in steatohepatitis: its role in Mallory-Denk body formation. Experimental and molecular pathology 90: 252–256, 2011.

37. French SW, French BA, Oliva J, Li J, Bardag-Gorce F, Tillman B, and Canaan A. FAT10 knock out mice livers fail to develop Mallory-Denk bodies in the DDC mouse model. Experimental and molecular pathology 93: 309–314, 2012.

38. Fromm H, Carlson GL, Hofmann AF, Farivar S, and Amin P. Metabolism in man of 7-ketolithocholic acid: precursor of cheno- and ursodeoxycholic acids. The American journal of physiology 239: G161–166, 1980.

39. Gautier-Stein A, Soty M, Chilloux J, Zitoun C, Rajas F, and Mithieux G. Glucotoxicity induces glucose-6-phosphatase catalytic unit expression by acting on the interaction of HIF-1alpha with CREB-binding protein. Diabetes 61: 2451–2460, 2012.

40. Gehrke N, Hovelmeyer N, Waisman A, Straub BK, Weinmann-Menke J, Worns MA, Galle PR, and Schattenberg JM. Hepatocyte-specific deletion of IL1-RI attenuates liver injury by blocking IL-1 driven autoinflammation. Journal of hepatology 68: 986–995, 2018.

41. Goodwin B, Watson MA, Kim H, Miao J, Kemper JK, and Kliewer SA. Differential regulation of rat and human CYP7A1 by the nuclear oxysterol receptor liver X receptor-alpha. Mol Endocrinol 17: 386–394, 2003.

42. Granados-Silvestre MA, Ortiz-Lopez MG, Granados J, Canizales-Quinteros S, Penaloza-Espinosa RI, Lechuga C, Acuna-Alonzo V, Sanchez-Pozos K, and Menjivar M. Susceptibility background for type 2 diabetes in eleven Mexican Indigenous populations: HNF4A gene analysis. Molecular genetics and genomics: MGG 292: 1209–1219, 2017.

43. Gross DA, Zhan C, and Silver DL. Direct binding of triglyceride to fat storage-inducing transmembrane proteins 1 and 2 is important for lipid droplet formation. Proceedings of the National Academy of Sciences of the United States of America 108: 19581–19586, 2011.

44. Gu J, Weng Y, Zhang QY, Cui H, Behr M, Wu L, Yang W, Zhang L, and Ding X. Liver-specific deletion of the NADPH-cytochrome P450 reductase gene: impact on plasma cholesterol homeostasis and the function and regulation of microsomal cytochrome P450 and heme oxygenase. The Journal of biological chemistry 278: 25895–25901, 2003.

45. Hannah VC, Ou J, Luong A, Goldstein JL, and Brown MS. Unsaturated fatty acids down-regulate srebp isoforms 1a and 1c by two mechanisms in HEK-293 cells. The Journal of biological chemistry 276: 4365–4372, 2001.

46. Hata T, Rehman F, Hori T, and Nguyen JH. GABA, gamma-Aminobutyric Acid, Protects Against Severe Liver Injury. The Journal of surgical research 236: 172–183, 2019.

47. Hayhurst GP, Lee YH, Lambert G, Ward JM, and Gonzalez FJ. Hepatocyte nuclear factor 4alpha (nuclear receptor 2A1) is essential for maintenance of hepatic gene expression and lipid homeostasis. Molecular and cellular biology 21: 1393–1403, 2001.

48. Heuman DM. Quantitative estimation of the hydrophilic-hydrophobic balance of mixed bile salt solutions. Journal of lipid research 30: 719–730, 1989.

49. Hiroyama M, Aoyagi T, Fujiwara Y, Birumachi J, Shigematsu Y, Kiwaki K, Tasaki R, Endo F, and Tanoue A. Hypermetabolism of fat in V1a vasopressin receptor knockout mice. Mol Endocrinol 21: 247–258, 2007.

50. Honda A, Miyazaki T, Iwamoto J, Hirayama T, Morishita Y, Monma T, Ueda H, Mizuno S, Sugiyama F, Takahashi S, and Ikegami T. Regulation of bile acid metabolism in mouse models with hydrophobic bile acid composition. Journal of lipid research 61: 54–69, 2020.

51. Hotta Y, Nakamura H, Konishi M, Murata Y, Takagi H, Matsumura S, Inoue K, Fushiki T, and Itoh N. Fibroblast growth factor 21 regulates lipolysis in white adipose tissue but is not required for ketogenesis and triglyceride clearance in liver. Endocrinology 150: 4625–4633, 2009.

52. Huang KW, Reebye V, Czysz K, Ciriello S, Dorman S, Reccia I, Lai HS, Peng L, Kostomitsopoulos N, Nicholls J, Habib RS, Tomalia DA, Saetrom P, Wilkes E, Cutillas P, Rossi JJ, and Habib NA. Liver Activation of Hepatocellular Nuclear Factor-4alpha by Small Activating RNA Rescues Dyslipidemia and Improves Metabolic Profile. Mol Ther Nucleic Acids 19: 361–370, 2020.

53. Hwang-Verslues WW, and Sladek FM. Nuclear receptor hepatocyte nuclear factor 4alpha1 competes with oncoprotein c-Myc for control of the p21/WAF1 promoter. Mol Endocrinol 22: 78–90, 2008.

54. Inoue Y, Yu AM, Inoue J, and Gonzalez FJ. Hepatocyte nuclear factor 4alpha is a central regulator of bile acid conjugation. The Journal of biological chemistry 279: 2480–2489, 2004.

55. Iordanidou P, Aggelidou E, Demetriades C, and Hadzopoulou-Cladaras M. Distinct amino acid residues may be involved in coactivator and ligand interactions in hepatocyte nuclear factor-4alpha. The Journal of biological chemistry 280: 21810–21819, 2005.

56. Ishida H, Kuruta Y, Gotoh O, Yamashita C, Yoshida Y, and Noshiro M. Structure, evolution, and liver-specific expression of sterol 12alpha-hydroxylase P450 (CYP8B). Journal of biochemistry 126: 19–25, 1999.

57. John K, Marino JS, Sanchez ER, and Hinds TD, Jr. The glucocorticoid receptor: cause of or cure for obesity? American journal of physiology Endocrinology and metabolism 310: E249–257, 2016.

58. Kadereit B, Kumar P, Wang WJ, Miranda D, Snapp EL, Severina N, Torregroza I, Evans T, and Silver DL. Evolutionarily conserved gene family important for fat storage. Proceedings of the National Academy of Sciences of the United States of America 105: 94–99, 2008.

59. Kelley GL, and Azhar S. Reversal of high dietary fructose-induced PPARalpha suppression by oral administration of lipoxygenase/cyclooxygenase inhibitors. Nutrition & metabolism 2: 18, 2005.

60. Kim JK, Fillmore JJ, Chen Y, Yu C, Moore IK, Pypaert M, Lutz EP, Kako Y, Velez-Carrasco W, Goldberg IJ, Breslow JL, and Shulman GI. Tissue-specific overexpression of lipoprotein lipase causes tissue-specific insulin resistance. Proceedings of the National Academy of Sciences of the United States of America 98: 7522–7527, 2001.

61. Kimura H. Histone modifications for human epigenome analysis. J Hum Genet 58: 439–445, 2013.

62. Kohli R, Kirby M, Xanthakos SA, Softic S, Feldstein AE, Saxena V, Tang PH, Miles L, Miles MV, Balistreri WF, Woods SC, and Seeley RJ. High-fructose, medium chain trans fat diet induces liver fibrosis and elevates plasma coenzyme Q9 in a novel murine model of obesity and nonalcoholic steatohepatitis. Hepatology 52: 934–944, 2010.

63. Komati R, Spadoni D, Zheng S, Sridhar J, Riley KE, and Wang G. Ligands of Therapeutic Utility for the Liver X Receptors. Molecules 22: 2017.

64. Konopelska S, Kienitz T, Hughes B, Pirlich M, Bauditz J, Lochs H, Strasburger CJ, Stewart PM, and Quinkler M. Hepatic 11beta-HSD1 mRNA expression in fatty liver and nonalcoholic steatohepatitis. Clinical endocrinology 70: 554–560, 2009.

65. Kopec AK, Joshi N, Towery KL, Kassel KM, Sullivan BP, Flick MJ, and Luyendyk JP. Thrombin inhibition with dabigatran protects against high-fat diet-induced fatty liver disease in mice. The Journal of pharmacology and experimental therapeutics 351: 288–297, 2014.

66. Krakowiak PA, Wassif CA, Kratz L, Cozma D, Kovarova M, Harris G, Grinberg A, Yang Y, Hunter AG, Tsokos M, Kelley RI, and Porter FD. Lathosterolosis: an inborn error of human and murine cholesterol synthesis due to lathosterol 5-desaturase deficiency. Human molecular genetics 12: 1631–1641, 2003.

67. Kronenberg F, Stuhlinger M, Trenkwalder E, Geethanjali FS, Pachinger O, von Eckardstein A, and Dieplinger H. Low apolipoprotein A-IV plasma concentrations in men with coronary artery disease. J Am Coll Cardiol 36: 751–757, 2000.

68. Langlais D, Couture C, Balsalobre A, and Drouin J. The Stat3/GR interaction code: predictive value of direct/indirect DNA recruitment for transcription outcome. Molecular cell 47: 38–49, 2012.

69. Lee YK, Dell H, Dowhan DH, Hadzopoulou-Cladaras M, and Moore DD. The orphan nuclear receptor SHP inhibits hepatocyte nuclear factor 4 and retinoid X receptor transactivation: two mechanisms for repression. Molecular and cellular biology 20: 187–195, 2000.

70. Lemke U, Krones-Herzig A, Berriel Diaz M, Narvekar P, Ziegler A, Vegiopoulos A, Cato AC, Bohl S, Klingmuller U, Screaton RA, Muller-Decker K, Kersten S, and Herzig S. The glucocorticoid receptor controls hepatic dyslipidemia through Hes1. Cell metabolism 8: 212–223, 2008.

71. Li T, and Apte U. Bile Acid Metabolism and Signaling in Cholestasis, Inflammation, and Cancer. Adv Pharmacol 74: 263–302, 2015.

72. Li T, and Chiang JY. Rifampicin induction of CYP3A4 requires pregnane X receptor cross talk with hepatocyte nuclear factor 4alpha and coactivators, and suppression of small heterodimer partner gene expression. Drug metabolism and disposition: the biological fate of chemicals 34: 756–764, 2006.

73. Li T, Jahan A, and Chiang JY. Bile acids and cytokines inhibit the human cholesterol 7 alpha-hydroxylase gene via the JNK/c-jun pathway in human liver cells. Hepatology 43: 1202–1210, 2006.

74. Li T, Owsley E, Matozel M, Hsu P, Novak CM, and Chiang JY. Transgenic expression of cholesterol 7alpha-hydroxylase in the liver prevents high-fat diet-induced obesity and insulin resistance in mice. Hepatology 52: 678–690, 2010.

75. Lim S, Taskinen MR, and Boren J. Crosstalk between nonalcoholic fatty liver disease and cardiometabolic syndrome. Obesity reviews: an official journal of the International Association for the Study of Obesity 2018.

76. Liu H, Li J, Tillman B, French BA, and French SW. Ufmylation and FATylation pathways are downregulated in human alcoholic and nonalcoholic steatohepatitis, and mice fed DDC, where Mallory-Denk bodies (MDBs) form. Experimental and molecular pathology 97: 81–88, 2014.

77. Longo M, Crosignani A, and Podda M. Hyperlipidemia in Chronic Cholestatic Liver Disease. Curr Treat Options Gastroenterol 4: 111–114, 2001.

78. Longuet C, Robledo AM, Dean ED, Dai C, Ali S, McGuinness I, de Chavez V, Vuguin PM, Charron MJ, Powers AC, and Drucker DJ. Liver-specific disruption of the murine glucagon receptor produces alpha-cell hyperplasia: evidence for a circulating alpha-cell growth factor. Diabetes 62: 1196–1205, 2013.

79. Lu H. Crosstalk of 51-Monophosphate-Activated Protein Kinase (AMPK) with Extracellular and Intracellular Signaling Pathways in the Regulation of Nutrient Metabolism and Cell Survival in the Liver. Current Pharmacology Reports 3: 162–175, 2017.

80. Lu H. Crosstalk of HNF4alpha with extracellular and intracellular signaling pathways in the regulation of hepatic metabolism of drugs and lipids. Acta Pharm Sin B 6: 393–408, 2016.

81. Lu H, Cui W, and Klaassen CD. Nrf2 protects against 2,3,7,8-tetrachlorodibenzo-p-dioxin (TCDD)-induced oxidative injury and steatohepatitis. Toxicology and applied pharmacology 256: 122–135, 2011.

82. Lu H, Gonzalez FJ, and Klaassen C. Alterations in hepatic mRNA expression of phase II enzymes and xenobiotic transporters after targeted disruption of hepatocyte nuclear factor 4 alpha. Toxicological sciences: an official journal of the Society of Toxicology 118: 380–390, 2010.

83. Magee N, Zou A, Ghosh P, Ahamed F, Delker D, and Zhang Y. Disruption of hepatic small heterodimer partner induces dissociation of steatosis and inflammation in experimental nonalcoholic steatohepatitis. The Journal of biological chemistry 295: 994–1008, 2020.

84. Melander O. Vasopressin, from Regulator to Disease Predictor for Diabetes and Cardiometabolic Risk. Annals of nutrition & metabolism 68 Suppl 2: 24–28, 2016.

85. Meng J, Feng M, Dong W, Zhu Y, Li Y, Zhang P, Wu L, Li M, Lu Y, Chen H, Liu X, Sun H, and Tong X. Identification of HNF-4alpha as a key transcription factor to promote ChREBP expression in response to glucose. Scientific reports 6: 23944, 2016.

86. Michailidou Z, Carter RN, Marshall E, Sutherland HG, Brownstein DG, Owen E, Cockett K, Kelly V, Ramage L, Al-Dujaili EA, Ross M, Maraki I, Newton K, Holmes MC, Seckl JR, Morton NM, Kenyon CJ, and Chapman KE. Glucocorticoid receptor haploinsufficiency causes hypertension and attenuates hypothalamic-pituitary-adrenal axis and blood pressure adaptions to high-fat diet. FASEB journal: official publication of the Federation of American Societies for Experimental Biology 22: 3896–3907, 2008.

87. Mittelstadt PR, Monteiro JP, and Ashwell JD. Thymocyte responsiveness to endogenous glucocorticoids is required for immunological fitness. The Journal of clinical investigation 122: 2384–2394, 2012.

88. Montagner A, Polizzi A, Fouche E, Ducheix S, Lippi Y, Lasserre F, Barquissau V, Regnier M, Lukowicz C, Benhamed F, Iroz A, Bertrand-Michel J, Al Saati T, Cano P, Mselli-Lakhal L, Mithieux G, Rajas F, Lagarrigue S, Pineau T, Loiseau N, Postic C, Langin D, Wahli W, and Guillou H. Liver PPARalpha is crucial for whole-body fatty acid homeostasis and is protective against NAFLD. Gut 65: 1202–1214, 2016.

89. Monte MJ, Marin JJ, Antelo A, and Vazquez-Tato J. Bile acids: chemistry, physiology, and pathophysiology. World journal of gastroenterology 15: 804–816, 2009.

90. Mork LM, Strom SC, Mode A, and Ellis EC. Addition of Dexamethasone Alters the Bile Acid Composition by Inducing CYP8B1 in Primary Cultures of Human Hepatocytes. Journal of clinical and experimental hepatology 6: 87–93, 2016.

91. Mueller KM, Kornfeld JW, Friedbichler K, Blaas L, Egger G, Esterbauer H, Hasselblatt P, Schlederer M, Haindl S, Wagner KU, Engblom D, Haemmerle G, Kratky D, Sexl V, Kenner L, Kozlov AV, Terracciano L, Zechner R, Schuetz G, Casanova E, Pospisilik JA, Heim MH, and Moriggl R. Impairment of hepatic growth hormone and glucocorticoid receptor signaling causes steatosis and hepatocellular carcinoma in mice. Hepatology 54: 1398–1409, 2011.

92. Mutch DM, Klocke B, Morrison P, Murray CA, Henderson CJ, Seifert M, and Williamson G. The disruption of hepatic cytochrome p450 reductase alters mouse lipid metabolism. Journal of proteome research 6: 3976–3984, 2007.

93. Nader N, Ng SS, Lambrou GI, Pervanidou P, Wang Y, Chrousos GP, and Kino T. AMPK regulates metabolic actions of glucocorticoids by phosphorylating the glucocorticoid receptor through p38 MAPK. Mol Endocrinol 24: 1748–1764, 2010.

94. Najt CP, Senthivinayagam S, Aljazi MB, Fader KA, Olenic SD, Brock JR, Lydic TA, Jones AD, and Atshaves BP. Liver-specific loss of Perilipin 2 alleviates diet-induced hepatic steatosis, inflammation, and fibrosis. American journal of physiology Gastrointestinal and liver physiology 310: G726–738, 2016.

95. Nguyen LN, Ma D, Shui G, Wong P, Cazenave-Gassiot A, Zhang X, Wenk MR, Goh EL, and Silver DL. Mfsd2a is a transporter for the essential omega-3 fatty acid docosahexaenoic acid. Nature 509: 503–506, 2014.

96. Niemi NM, Wilson GM, Overmyer KA, Vogtle FN, Myketin L, Lohman DC, Schueler KL, Attie AD, Meisinger C, Coon JJ, and Pagliarini DJ. Pptc7 is an essential phosphatase for promoting mammalian mitochondrial metabolism and biogenesis. Nature communications 10: 3197, 2019.

97. Ochoa-Sanchez R, and Rose CF. Pathogenesis of Hepatic Encephalopathy in Chronic Liver Disease. Journal of clinical and experimental hepatology 8: 262–271, 2018.

98. Oliveras-Ferraros C, Vazquez-Martin A, Fernandez-Real JM, and Menendez JA. AMPK-sensed cellular energy state regulates the release of extracellular Fatty Acid Synthase. Biochemical and biophysical research communications 378: 488–493, 2009.

99. Olivieri O, Bassi A, Stranieri C, Trabetti E, Martinelli N, Pizzolo F, Girelli D, Friso S, Pignatti PF, and Corrocher R. Apolipoprotein C-III, metabolic syndrome, and risk of coronary artery disease. Journal of lipid research 44: 2374–2381, 2003.

100. Onica T, Nichols K, Larin M, Ng L, Maslen A, Dvorak Z, Pascussi JM, Vilarem MJ, Maurel P, and Kirby GM. Dexamethasone-mediated up-regulation of human CYP2A6 involves the glucocorticoid receptor and increased binding of hepatic nuclear factor 4 alpha to the proximal promoter. Molecular pharmacology 73: 451–460, 2008.

101. Oosterveer MH, Grefhorst A, van Dijk TH, Havinga R, Staels B, Kuipers F, Groen AK, and Reijngoud DJ. Fenofibrate simultaneously induces hepatic fatty acid oxidation, synthesis, and elongation in mice. The Journal of biological chemistry 284: 34036–34044, 2009.

102. Patel R, Bookout AL, Magomedova L, Owen BM, Consiglio GP, Shimizu M, Zhang Y, Mangelsdorf DJ, Kliewer SA, and Cummins CL. Glucocorticoids regulate the metabolic hormone FGF21 in a feed-forward loop. Mol Endocrinol 29: 213–223, 2015.

103. Paterson JM, Morton NM, Fievet C, Kenyon CJ, Holmes MC, Staels B, Seckl JR, and Mullins JJ. Metabolic syndrome without obesity: Hepatic overexpression of 11beta-hydroxysteroid dehydrogenase type 1 in transgenic mice. Proceedings of the National Academy of Sciences of the United States of America 101: 7088–7093, 2004.

104. Plump AS, Azrolan N, Odaka H, Wu L, Jiang X, Tall A, Eisenberg S, and Breslow JL. ApoA-I knockout mice: characterization of HDL metabolism in homozygotes and identification of a post-RNA mechanism of apoA-I up-regulation in heterozygotes. Journal of lipid research 38: 1033–1047, 1997.

105. Privitera G, Spadaro L, Marchisello S, Fede G, and Purrello F. Abnormalities of Lipoprotein Levels in Liver Cirrhosis: Clinical Relevance. Digestive diseases and sciences 63: 16–26, 2018.

106. Pullinger CR, Eng C, Salen G, Shefer S, Batta AK, Erickson SK, Verhagen A, Rivera CR, Mulvihill SJ, Malloy MJ, and Kane JP. Human cholesterol 7alpha-hydroxylase (CYP7A1) deficiency has a hypercholesterolemic phenotype. The Journal of clinical investigation 110: 109–117, 2002.

107. Qu X, Lam E, Doughman YQ, Chen Y, Chou YT, Lam M, Turakhia M, Dunwoodie SL, Watanabe M, Xu B, Duncan SA, and Yang YC. Cited2, a coactivator of HNF4alpha, is essential for liver development. The EMBO journal 26: 4445–4456, 2007.

108. Ratman D, Mylka V, Bougarne N, Pawlak M, Caron S, Hennuyer N, Paumelle R, De Cauwer L, Thommis J, Rider MH, Libert C, Lievens S, Tavernier J, Staels B, and De Bosscher K. Chromatin recruitment of activated AMPK drives fasting response genes co-controlled by GR and PPARalpha. Nucleic acids research 44: 10539–10553, 2016.

109. Reid BN, Ables GP, Otlivanchik OA, Schoiswohl G, Zechner R, Blaner WS, Goldberg IJ, Schwabe RF, Chua SC, Jr., and Huang LS. Hepatic overexpression of hormone-sensitive lipase and adipose triglyceride lipase promotes fatty acid oxidation, stimulates direct release of free fatty acids, and ameliorates steatosis. The Journal of biological chemistry 283: 13087–13099, 2008.

110. Repa JJ, Liang G, Ou J, Bashmakov Y, Lobaccaro JM, Shimomura I, Shan B, Brown MS, Goldstein JL, and Mangelsdorf DJ. Regulation of mouse sterol regulatory element-binding protein-1c gene (SREBP-1c) by oxysterol receptors, LXRalpha and LXRbeta. Genes & development 14: 2819–2830, 2000.

111. Rhee J, Inoue Y, Yoon JC, Puigserver P, Fan M, Gonzalez FJ, and Spiegelman BM. Regulation of hepatic fasting response by PPARgamma coactivator-1alpha (PGC-1): requirement for hepatocyte nuclear factor 4alpha in gluconeogenesis. Proceedings of the National Academy of Sciences of the United States of America 100: 4012–4017, 2003.

112. Robinson JT, Thorvaldsdottir H, Winckler W, Guttman M, Lander ES, Getz G, and Mesirov JP. Integrative genomics viewer. Nature biotechnology 29: 24–26, 2011.

113. Roqueta-Rivera M, Esquejo RM, Phelan PE, Sandor K, Daniel B, Foufelle F, Ding J, Li X, Khorasanizadeh S, and Osborne TF. SETDB2 Links Glucocorticoid to Lipid Metabolism through Insig2a Regulation. Cell metabolism 24: 474–484, 2016.

114. Sato M, Kawakami T, Kondoh M, Takiguchi M, Kadota Y, Himeno S, and Suzuki S. Development of high-fat-diet-induced obesity in female metallothionein-null mice. FASEB J 24: 2375–2384, 2010.

115. Sato O, Kuriki C, Fukui Y, and Motojima K. Dual promoter structure of mouse and human fatty acid translocase/CD36 genes and unique transcriptional activation by peroxisome proliferator-activated receptor alpha and gamma ligands. The Journal of biological chemistry 277: 15703–15711, 2002.

116. Sayin SI, Wahlstrom A, Felin J, Jantti S, Marschall HU, Bamberg K, Angelin B, Hyotylainen T, Oresic M, and Backhed F. Gut microbiota regulates bile acid metabolism by reducing the levels of tauro-beta-muricholic acid, a naturally occurring FXR antagonist. Cell metabolism 17: 225–235, 2013.

117. Schregle R, Mah MM, Mueller S, Aichem A, Basler M, and Groettrup M. The expression profile of the ubiquitin-like modifier FAT10 in immune cells suggests cell type-specific functions. Immunogenetics 70: 429–438, 2018.

118. Schuler M, Dierich A, Chambon P, and Metzger D. Efficient temporally controlled targeted somatic mutagenesis in hepatocytes of the mouse. Genesis 39: 167–172, 2004.

119. Shao YX, Huang M, Cui W, Feng LJ, Wu Y, Cai Y, Li Z, Zhu X, Liu P, Wan Y, Ke H, and Luo HB. Discovery of a phosphodiesterase 9A inhibitor as a potential hypoglycemic agent. J Med Chem 57: 10304–10313, 2014.

120. She P, Shiota M, Shelton KD, Chalkley R, Postic C, and Magnuson MA. Phosphoenolpyruvate carboxykinase is necessary for the integration of hepatic energy metabolism. Molecular and cellular biology 20: 6508–6517, 2000.

121. Sheng L, Cho KW, Zhou Y, Shen H, and Rui L. Lipocalin 13 protein protects against hepatic steatosis by both inhibiting lipogenesis and stimulating fatty acid beta-oxidation. The Journal of biological chemistry 286: 38128–38135, 2011.

122. Shi X, Shi W, Li Q, Song B, Wan M, Bai S, and Cao X. A glucocorticoid-induced leucine-zipper protein, GILZ, inhibits adipogenesis of mesenchymal cells. EMBO reports 4: 374–380, 2003.

123. Shimizu N, Maruyama T, Yoshikawa N, Matsumiya R, Ma Y, Ito N, Tasaka Y, Kuribara-Souta A, Miyata K, Oike Y, Berger S, Schutz G, Takeda S, and Tanaka H. A muscle-liver-fat signalling axis is essential for central control of adaptive adipose remodelling. Nature communications 6: 6693, 2015.

124. Shin MJ, and Krauss RM. Apolipoprotein CIII bound to apoB-containing lipoproteins is associated with small, dense LDL independent of plasma triglyceride levels in healthy men. Atherosclerosis 211: 337–341, 2010.

125. Song C, Hiipakka RA, and Liao S. Selective activation of liver X receptor alpha by 6alpha-hydroxy bile acids and analogs. Steroids 65: 423–427, 2000.

126. Stafford JM, Wilkinson JC, Beechem JM, and Granner DK. Accessory factors facilitate the binding of glucocorticoid receptor to the phosphoenolpyruvate carboxykinase gene promoter. The Journal of biological chemistry 276: 39885–39891, 2001.

127. Su Q, Baker C, Christian P, Naples M, Tong X, Zhang K, Santha M, and Adeli K. Hepatic mitochondrial and ER stress induced by defective PPARalpha signaling in the pathogenesis of hepatic steatosis. American journal of physiology Endocrinology and metabolism 306: E1264–1273, 2014.

128. Takahashi S, Fukami T, Masuo Y, Brocker CN, Xie C, Krausz KW, Wolf CR, Henderson CJ, and Gonzalez FJ. Cyp2c70 is responsible for the species difference in bile acid metabolism between mice and humans. Journal of lipid research 57: 2130–2137, 2016.

129. Takiguchi S, Ayaori M, Yakushiji E, Nishida T, Nakaya K, Sasaki M, Iizuka M, Uto-Kondo H, Terao Y, Yogo M, Komatsu T, Ogura M, and Ikewaki K. Hepatic Overexpression of Endothelial Lipase Lowers High-Density Lipoprotein but Maintains Reverse Cholesterol Transport in Mice: Role of Scavenger Receptor Class B Type I/ATP-Binding Cassette Transporter A1-Dependent Pathways. Arteriosclerosis, thrombosis, and vascular biology 38: 1454–1467, 2018.

130. Tao H, Aakula S, Abumrad NN, and Hajri T. Peroxisome proliferator-activated receptor-gamma regulates the expression and function of very-low-density lipoprotein receptor. American journal of physiology Endocrinology and metabolism 298: E68–79, 2010.

131. Targett-Adams P, McElwee MJ, Ehrenborg E, Gustafsson MC, Palmer CN, and McLauchlan J. A PPAR response element regulates transcription of the gene for human adipose differentiation-related protein. Biochimica et biophysica acta 1728: 95–104, 2005.

132. Tchorz JS, Kinter J, Muller M, Tornillo L, Heim MH, and Bettler B. Notch2 signaling promotes biliary epithelial cell fate specification and tubulogenesis during bile duct development in mice. Hepatology 50: 871–879, 2009.

133. Thakare R, Alamoudi JA, Gautam N, Rodrigues AD, and Alnouti Y. Species differences in bile acids I. Plasma and urine bile acid composition. J Appl Toxicol 38: 1323–1335, 2018.

134. Thomas AM, Hart SN, Li G, Lu H, Fang Y, Fang J, Zhong XB, and Guo GL. Hepatocyte nuclear factor 4 alpha and farnesoid X receptor co-regulates gene transcription in mouse livers on a genome-wide scale. Pharm Res 30: 2188–2198, 2013.

135. Trefflich I, Marschall HU, Giuseppe RD, Stahlman M, Michalsen A, Lampen A, Abraham K, and Weikert C. Associations between Dietary Patterns and Bile Acids-Results from a Cross-Sectional Study in Vegans and Omnivores. Nutrients 12: 2019.

136. Vandevyver S, Dejager L, and Libert C. Comprehensive overview of the structure and regulation of the glucocorticoid receptor. Endocrine reviews 35: 671–693, 2014.

137. VerHague MA, Cheng D, Weinberg RB, and Shelness GS. Apolipoprotein A-IV expression in mouse liver enhances triglyceride secretion and reduces hepatic lipid content by promoting very low density lipoprotein particle expansion. Arteriosclerosis, thrombosis, and vascular biology 33: 2501–2508, 2013.

138. Waller-Evans H, Hue C, Fearnside J, Rothwell AR, Lockstone HE, Calderari S, Wilder SP, Cazier JB, Scott J, and Gauguier D. Nutrigenomics of high fat diet induced obesity in mice suggests relationships between susceptibility to fatty liver disease and the proteasome. PLoS ONE 8: e82825, 2013.

139. Wang H, Chen J, Hollister K, Sowers LC, and Forman BM. Endogenous bile acids are ligands for the nuclear receptor FXR/BAR. Molecular cell 3: 543–553, 1999.

140. Watanabe M, Houten SM, Wang L, Moschetta A, Mangelsdorf DJ, Heyman RA, Moore DD, and Auwerx J. Bile acids lower triglyceride levels via a pathway involving FXR, SHP, and SREBP-1c. The Journal of clinical investigation 113: 1408–1418, 2004.

141. Weng W, and Breslow JL. Dramatically decreased high density lipoprotein cholesterol, increased remnant clearance, and insulin hypersensitivity in apolipoprotein A-II knockout mice suggest a complex role for apolipoprotein A-II in atherosclerosis susceptibility. Proceedings of the National Academy of Sciences of the United States of America 93: 14788–14794, 1996.

142. Wilson CG, Tran JL, Erion DM, Vera NB, Febbraio M, and Weiss EJ. Hepatocyte-Specific Disruption of CD36 Attenuates Fatty Liver and Improves Insulin Sensitivity in HFD-Fed Mice. Endocrinology 157: 570–585, 2016.

143. Wong S, Tan K, Carey KT, Fukushima A, Tiganis T, and Cole TJ. Glucocorticoids stimulate hepatic and renal catecholamine inactivation by direct rapid induction of the dopamine sulfotransferase Sult1d1. Endocrinology 151: 185–194, 2010.

144. Wortham M, Czerwinski M, He L, Parkinson A, and Wan YJ. Expression of constitutive androstane receptor, hepatic nuclear factor 4 alpha, and P450 oxidoreductase genes determines interindividual variability in basal expression and activity of a broad scope of xenobiotic metabolism genes in the human liver. Drug metabolism and disposition: the biological fate of chemicals 35: 1700–1710, 2007.

145. Xie X, Liao H, Dang H, Pang W, Guan Y, Wang X, Shyy JY, Zhu Y, and Sladek FM. Down-regulation of hepatic HNF4alpha gene expression during hyperinsulinemia via SREBPs. Mol Endocrinol 23: 434–443, 2009.

146. Xu Y, Zalzala M, Xu J, Li Y, Yin L, and Zhang Y. A metabolic stress-inducible miR-34a-HNF4alpha pathway regulates lipid and lipoprotein metabolism. Nature communications 6: 7466, 2015.

147. Yan F, Wang Q, Xu C, Cao M, Zhou X, Wang T, Yu C, Jing F, Chen W, Gao L, and Zhao J. Peroxisome proliferator-activated receptor alpha activation induces hepatic steatosis, suggesting an adverse effect. PloS one 9: e99245, 2014.

148. Yoshitsugu R, Kikuchi K, Iwaya H, Fujii N, Hori S, Lee DG, and Ishizuka S. Alteration of Bile Acid Metabolism by a High-Fat Diet Is Associated with Plasma Transaminase Activities and Glucose Intolerance in Rats. J Nutr Sci Vitaminol (Tokyo) 65: 45–51, 2019.

149. Yu KC, David C, Kadambi S, Stahl A, Hirata K, Ishida T, Quertermous T, Cooper AD, and Choi SY. Endothelial lipase is synthesized by hepatic and aorta endothelial cells and its expression is altered in apoE-deficient mice. Journal of lipid research 45: 1614–1623, 2004.

150. Zhai X, Yan K, Fan J, Niu M, Zhou Q, Zhou Y, and Chen H. The beta-catenin pathway contributes to the effects of leptin on SREBP-1c expression in rat hepatic stellate cells and liver fibrosis. British journal of pharmacology 169: 197–212, 2013.

151. Zhang M, Zhao Y, Li Z, and Wang C. Pyruvate dehydrogenase kinase 4 mediates lipogenesis and contributes to the pathogenesis of nonalcoholic steatohepatitis. Biochemical and biophysical research communications 495: 582–586, 2018.

152. Zhang Y, Hagedorn CH, and Wang L. Role of nuclear receptor SHP in metabolism and cancer. Biochimica et biophysica acta 1812: 893–908, 2011.

153. Zhang Y, Li D, and Sun B. Do housekeeping genes exist? PloS one 10: e0123691, 2015.

154. Zhang Y, and Wang L. Characterization of the mitochondrial localization of the nuclear receptor SHP and regulation of its subcellular distribution by interaction with Bcl2 and HNF4alpha. PloS one 8: e68491, 2013.

155. Zhou Y, and Rui L. Lipocalin 13 regulation of glucose and lipid metabolism in obesity. Vitamins and hormones 91: 369–383, 2013.

156. Zhu P, Wang Y, Du Y, He L, Huang G, Zhang G, Yan X, and Fan Z. C8orf4 negatively regulates self-renewal of liver cancer stem cells via suppression of NOTCH2 signalling. Nature communications 6: 7122, 2015.

157. Zhu Y, Zhao S, Deng Y, Gordillo R, Ghaben AL, Shao M, Zhang F, Xu P, Li Y, Cao H, Zagnitko O, Scott DA, Gupta RK, Xing C, Zhang BB, Lin HV, and Scherer PE. Hepatic GALE Regulates Whole-Body Glucose Homeostasis by Modulating Tff3 Expression. Diabetes 66: 2789–2799, 2017.

158. Zou X, Ramachandran P, Kendall TJ, Pellicoro A, Dora E, Aucott RL, Manwani K, Man TY, Chapman KE, Henderson NC, Forbes SJ, Webster SP, Iredale JP, Walker BR, and Michailidou Z. 11Beta-hydroxysteroid dehydrogenase-1 deficiency or inhibition enhances hepatic myofibroblast activation in murine liver fibrosis. Hepatology 67: 2167–2181, 2018.

